# IL-1 instructs para-bronchial cuff fibroblasts to organize lung antibody secreting cell niches during continued antigen exposure

**DOI:** 10.64898/2026.01.09.698642

**Authors:** Inés Lammens, Jozefien Declercq, Andrew S Brown, Ursula Smole, Kim Deswarte, Esther Hoste, Karen Herreman, Chananchida Sang-aram, Robin Browaeys, Antonio P Baptista, Ruth Seurinck, Yvan Saeys, Philippe Gevaert, Hamida Hammad, Bart N Lambrecht, Stijn Vanhee

## Abstract

Secondary lymphoid organs (SLO) are prototypic sites of antibody production, yet mucosal sites also generate and maintain mucosal antibodies during continued inflammation. Using a murine model of house dust mite–induced airway inflammation, we show that prolonged allergen exposure induces tertiary lymphoid organs (TLO) within para-bronchial adventitial cuffs. Within these regions, IL-1 signaling instructs fibroblasts to form niches that recruit, retain, and sustain lung antibody-secreting cells (ASCs) through chemokine induction. These mucosal ASCs share immunoglobulin repertoires with TLO-derived germinal center B cells. Thus, continued allergen exposure reshapes the lung microenvironment by converting para-bronchial fibroblasts into IL-1–dependent supportive niches for non-IgE antibody production, revealing a fibroblast-mediated mechanism for local immune regulation in chronic inflammation.

**One sentence summary:** Continued allergen exposure drives lung TLO to generate ASCs that home to IL1-instructed para-bronchial cuffs.

## INTRODUCTION

Continued inflammation in the airways typically elicits the induction of mucosal antibodies (*1, 2*). Besides viral infections, the airways are often enriched for non-IgE antibodies in the context of chronic allergic airway diseases (*3, 4*). While the role of systemic IgE in allergy is well established, far less is known about the induction and maintenance of these mucosal antibodies, particularly IgM, IgG and IgA isotypes.

Antibody production typically arises from cognate interactions between antigen-specific T and B cells within secondary lymphoid organs (SLO), such as lung-draining lymph nodes, where germinal centers (GCs) mediate somatic hypermutation (SHM), affinity maturation, and class switch recombination (CSR) (*5, 6*). GC-derived B cells differentiate into long-lived circulating and tissue resident memory B cells and antibody-secreting cells (ASCs) that populate bone marrow and medullary cords, maintaining systemic antibody levels (*7–11*). Beyond SLO, persistent antigen exposure can trigger ectopic GC formation within tertiary lymphoid organs (TLO) across many tissues (*12–14*). In models of respiratory viral infection, lung TLO generate ASCs that produce locally protective antibodies detectable in bronchoalveolar lavage fluid (*1, 2, 15–24*). From an evolutionary perspective, TLO even predate SLO, suggesting that inflammation-induced TLO represent an ancient blueprint for adaptive immunity (*25–27*). Their conservation and ability to arise across diverse tissues underscore a central role in regulating local immune defense and inflammation.

Within SLO, stromal cells tightly orchestrate GC and ASC responses (*28–32*), yet how analogous processes are organized in non-lymphoid tissues remains poorly understood. Notably, lung TLO consistently emerge in subepithelial layer surrounding conducting airways after chronic stimulation, implicating resident fibroblasts as potential structural and functional analogs of lymphoid stromal cells (*28*). Emerging evidence of fibroblast heterogeneity and immune-instructive function supports this concept (*33–35*), but how fibroblast subsets control mucosal antibody production remains unresolved.

Here, using a model of chronic house dust mite (HDM)–induced airway inflammation, we sought to uncover the origin and function of locally produced mucosal antibodies. We identify TLO induction as a key feature of prolonged allergen exposure, with TLO forming preferentially in the adventitial cuffs surrounding bronchovascular bundles. VDJ repertoire analyses revealed that lung ASCs arise from mature GC responses within these TLO. Spatial and single-cell transcriptomic profiling further showed that ASCs migrate from TLO into adjacent para-bronchial cuffs, where IL-1 instructs local fibroblasts to produce chemokines and cytokines supporting ASC retention. Thus, IL-1–responsive fibroblasts form specialized niches that link chronic inflammation to local antibody production, revealing a previously unrecognized stromal mechanism shaping mucosal immunity.

## RESULTS

### Continued HDM exposure induces mucosal antibodies

Although multiple antibody isotypes are present in the airways following continued inflammation, their origin and maintenance remains unclear. To dissect these mechanisms, we set up a prototypic chronic inflammation model using the house dust mite (HDM) extract model, exposing wild-type (WT) C57BL/6 mice to inhaled HDM for 5 weeks. We quantified immunoglobulin isotypes in serum and bronchoalveolar lavage (BAL) fluid 1 week after initial sensitization and at successive time points during repeated challenges (Fig. 1A). Prolonged HDM exposure induced a canonical antibody profile, with IgE, IgG1, IgA and IgM isotypes progressively increasing with continued exposure in both serum and BAL (Fig. 1B-E). In contrast, initial allergic airway inflammation declined with continued HDM exposure (fig. S1A-B).

**Figure 1:**
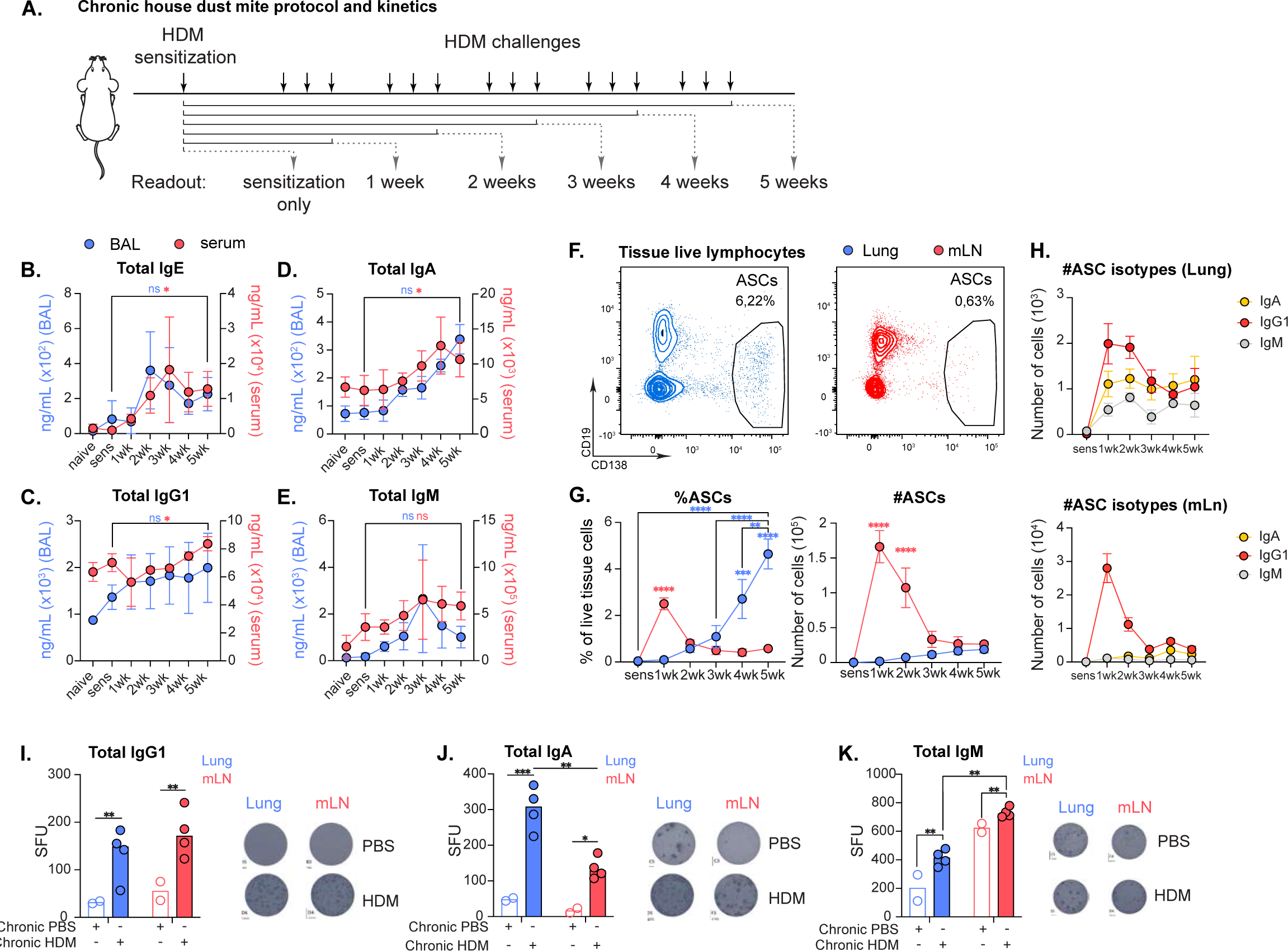
Chronic HDM exposure induces mucosal antibodies. A. Experimental set-up of prolonged HDM exposure model for kinetics analysis B. To E. Kinetics analysis of indicated immunoglobulin levels in BAL (blue, left y-axis) and serum (red, right y-axis) Total IgE kinetics C. Total IgG1 kinetics D. Total IgA kinetics E. Total IgM kinetics F. Flow cytometric analysis of lung (blue) and mLN (red) ASC kinetics. 5 week challenge timepoint shown G. Quantification of F H. Quantification of IgG1^+^, IgA^+^ and IgM^+^ ASC kinetics in lung and mLN I. ELISpot data from lung and mLN ASCs following chronic PBS and chronic HDM exposure For B-E, G-H data are shown as means + sem (n=4 for each timepoint). For I-K, bars represent median with each symbol representing 1 replicate (n=4 for each timepoint). P values are for 2-way-ANOVA (B-E, G-H) and Mann-Whitney U test (I-K) (*P<0,05, **P<0,01, ***P<0,001). Data are representative of 3 independent experiments.

We next examined whether the emergence of antibody responses coincided with the appearance of antibody-secreting cells (ASCs) in the lung mucosa and lung draining mediastinal lymph node (mLN) (fig. S1C). ASCs appeared in the mLN as early as 1 week after HDM exposure, but declined with continued challenge (Fig. 1F and G; fig. S1D). In contrast, lung ASCs accumulated progressively, albeit with slower kinetics and lower magnitude compared to the mLN (Fig. 1, F and G; fig. S1D). Early responses in both sites were dominated by IgG1⁺ ASCs, whereas prolonged HDM exposure led to comparable numbers of IgA⁺ and IgG1⁺ ASCs in the lung (Fig. 1H). In the mLN, IgG1⁺ ASCs remained predominant, with IgA⁺ ASCs emerging only at later time points, suggesting possible recirculation (Fig. 1H), which is in line with previous work (*36*). IgM⁺ ASCs followed similar kinetics in both compartments (Fig. 1H). ELISpot analysis confirmed comparable numbers of IgG1-secreting ASCs in lung and mLN (Fig. 1I). While IgM-secreting ASCs are increased in the mLN compared to the lung compartment, continued HDM exposure enriches the pulmonary mucosa with IgA-secreting ASCs (Fig. 1J and K). Together, these data suggest a compartmentalized ASC response upon continued inflammation, with an IgA-dominated response in the airway mucosa.

### Continued HDM exposure induces mature TLO germinal centers in the lung that generate mucosal ASCs

Because TLO can serve as local hubs for antibody production (*37–39*), we examined whether they form in lungs after continued HDM exposure. After 5 weeks, we observed well-organized TLO clusters composed of B cell follicles containing GL7⁺ activated B cells and dispersed T cells, consistent with GC–containing TLO (Fig. 2A). These structures harbored follicular dendritic cells (FDCs) within GC B cell zones and were organized around PNAd⁺ high endothelial venules (HEVs) and Lyve-1⁺ lymphatics (Fig. 2A and refs (*15, 16, 37, 39–44*)). Time-course analysis revealed that B cells initially accumulated in the adventitial cuff by 1 week of HDM exposure and progressively organized into clusters that matured into bona fide TLO by 4 weeks (fig. S2A). After 5 weeks, lung GC B and T follicular helper (Tfh) cells expressed *Bcl6,* yet lung Tfh to a lower extent compared to their mLN counterpart (Fig. 2B). Flow cytometry confirmed that GL7⁺CD95⁺ GC B cells appeared first in the mLN, followed by delayed TLO GC formation after 3 weeks (Fig. 2C; fig. S2, B-C and F). Whereas mLN GCs plateaued after 2 weeks, lung TLO GCs continued to expand (Fig. 2C). Given the concordant expression of GL7^+^CD95^+^ and *Bcl6^+^CD95^+^*, GL7 was used as a reliable GC marker for subsequent analyses (fig. S2, G-J). PD1⁺CXCR5⁺ Tfh cells followed similar kinetics, with robust accumulation in the lung by week 5, reflecting ongoing GC maturation (Fig. 2D; fig. S2, D, E, and H). Additionally, lung GC B cells expressed the class-switch enzyme Activation-induced cytidine deaminase (AID) in *AID*^CreERT2^ × *R26^TdTomato^* reporter mice (*45*) (Fig. 2, E and F) and segregated into light and dark zone subsets (fig. S3, A and B), confirming active GC function. Together, these data demonstrate that chronic HDM exposure drives the formation of mature, functional TLO GCs in the lung mucosa.

**Figure 2:**
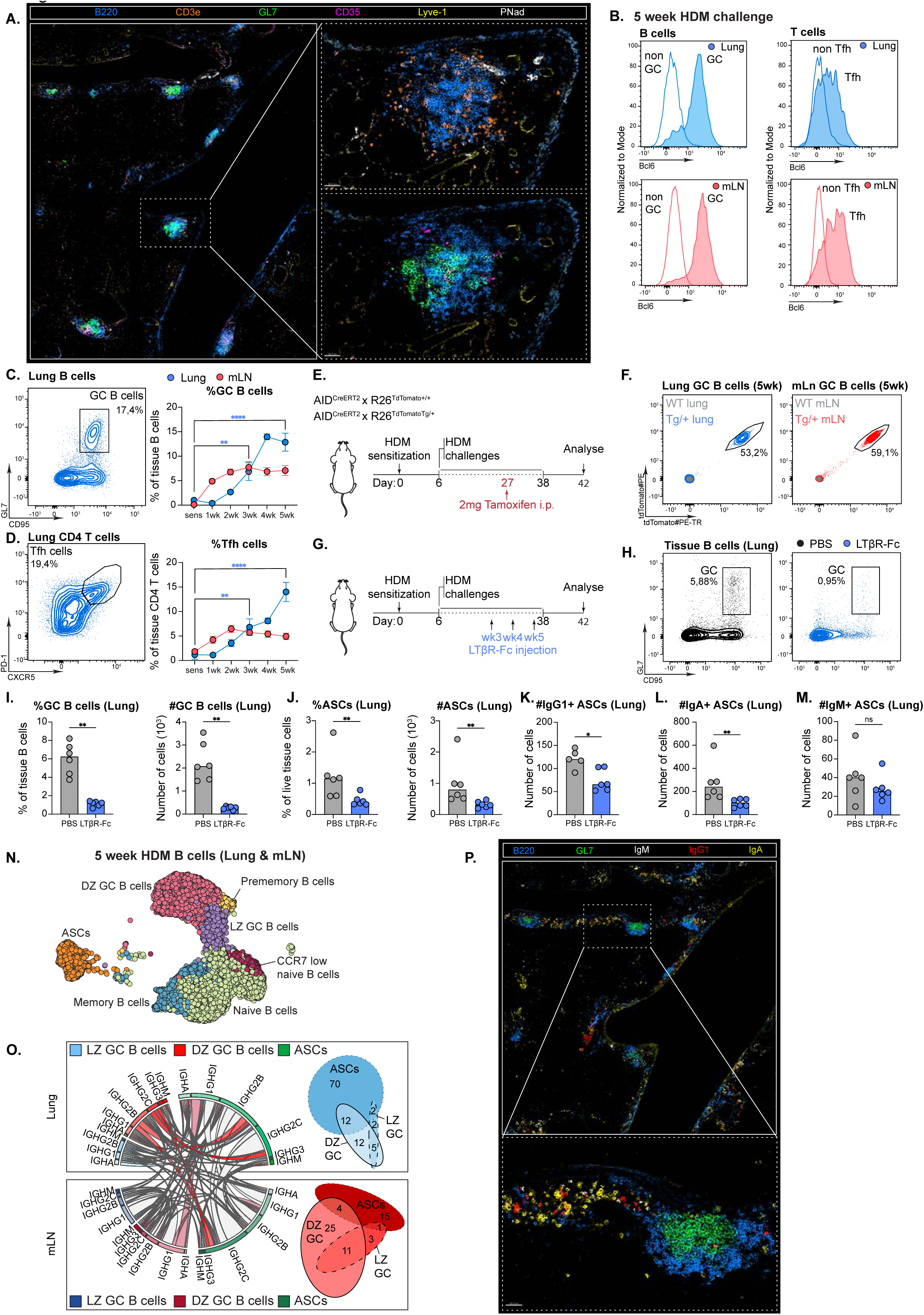
Continued HDM exposure induces mature TLO germinal centers in the lung that generate mucosal ASCs. A. Confocal imaging of frozen lung sections after 5 weeks of HDM challenges showing organized TLO consisting of B cell follicles (blue) that contain GCs (green) with FDCs (pink), and T cells (orange), lymphatics (yellow) and HEVs (white). Scale bars: 50μm B. Bcl6 expression in lung (blue) and mLN (red) Tfh cells (5 week timepoint) C. To D. Flow cytometric analysis and quantification of GC B cells (C) and D. T follicular helper cells (D) in lung (blue) and mLN (red) E. Experimental set-up for GC B cell labelling in AID^CreERT2^ x R26^TdTomato^ reporter mice F. Flow cytometric analysis of lung (blue) and mLN (red) GC B cell labelling in AID^CreERT2^ x R26^TdTomato^ reporter mice G. Experimental set-up for TLO ablation using LTβR-Fc H. Flow cytometric analysis of TLO GC ablation, showing mock (left, black) and LTβR-Fc (blue, right) I. Quantification of H J. Quantification of lung ASCs following mock and LTβR-Fc K. Quantification of lung IgG1^+^ ASCs following mock and LTβR-Fc L. Quantification of lung IgA^+^ ASCs following mock and LTβR-Fc M. Quantification of lung IgM^+^ ASCs following mock and LTβR-Fc N. Uniform manifold approximation and projection (UMAP) of integrated lung and mLN B cells following 5 weeks of HDM challenges O. BCR clonal relation in lung and mLN GC B cells and ASCs. Connections in white show relatedness between same phenotypic clusters between organs (GC-GC or ASC-ASC). Colored connections show relatedness between GC and ASCs. Quantification of unique clonal connections in the same organ shown in Venn diagrams. P. Confocal imaging of frozen lung sections after 5 weeks of HDM challenges, showing B cell follicles (blue) with GCs (green), and IgM+ (white), IgG1+ (red) and IgA+ (yellow) halos as a hallmark for ASCs. Scale bar: 50μm. For C-D, data are presented as means+sem (n=4 for each timepoint). For I-M, bars represent median with each symbol representing 1 replicate (n=6 for each timepoint). P values are for 2-way-ANOVA (C-D) and Mann-Whitney U test (I-M) (*P<0,05, **P<0,01, ***P<0,001). Data are representative of 3 independent experiments.

Given the coincident emergence of lung ASCs (Fig. 1F and G), we next examined whether TLO GCs contribute to the mucosal ASC pool, employing a previously published method to dissassemble TLO that depend on continuous signaling of lymphotoxin-β (*15*). Systemic administration of the LTβ antagonist LTβR-Fc from weeks 3 to 5 (Fig. 2G) effectively ablated lung TLO GC formation, reducing total ASCs as well as IgG1⁺ and IgA⁺ ASC numbers (Fig. 2, H–M; fig. S3C). mLN GC and ASC responses were less affected (fig. S3,D-J), suggesting that lung TLO GCs directly sustain mucosal ASC production, though a contribution from mLN GCs cannot be excluded.

To further uncover the relationship between TLO GCs and ASCs, we performed single-cell RNA, epitope, and immune receptor sequencing (CITE-R-seq) on sorted lung and mLN B cells (fig. S3K). Both compartments contained diverse B cell populations identified by canonical markers (Fig. 2N; fig. S3L). Despite the polyclonal settging, clonal analysis revealed BCR overlap between lung GC and ASC populations (14 shared clones), whereas mLN GCs primarily contributed to mLN ASCs (5 shared clones; Fig. 2O). These data further support lung TLO GCs as a local source of mucosa-resident ASCs. By week 5 of HDM exposure, these mucosal ASCs were enriched in TLO-adjacent niches, forming chain-like clusters suggestive of a local survival microenvironment (Fig. 2P). Collectively, continued HDM exposure promotes the formation of functional TLO GCs in the lung that give rise to mucosal ASCs residing in specialized niches in proximity to TLO.

### The para-bronchial cuff niche supports lung ASC residence

The adventitial cuff is a known site for tissue-resident immune cells and TLO formation (*15, 19, 46, 47*), yet how ASCs establish residence in the lung mucosa remains unclear. Our earlier findings suggested that distinct regions adjacent to the adventitial cuff support mucosal ASC localization (Fig. 2P). To define these niches, we examined the spatial kinetics of ASC distribution relative to lung architecture. As expected, TLO GCs developed within the adventitial cuff (Fig. 3A). Early in HDM exposure, ASCs were confined to this region; however, beginning at week 3, they progressively migrated toward subepithelial areas surrounding broncho-vascular bundles, which we termed the “para-bronchial cuff” (Fig. 3A). This redistribution became more pronounced with continued exposure (Fig. 3A).

**Figure 3:**
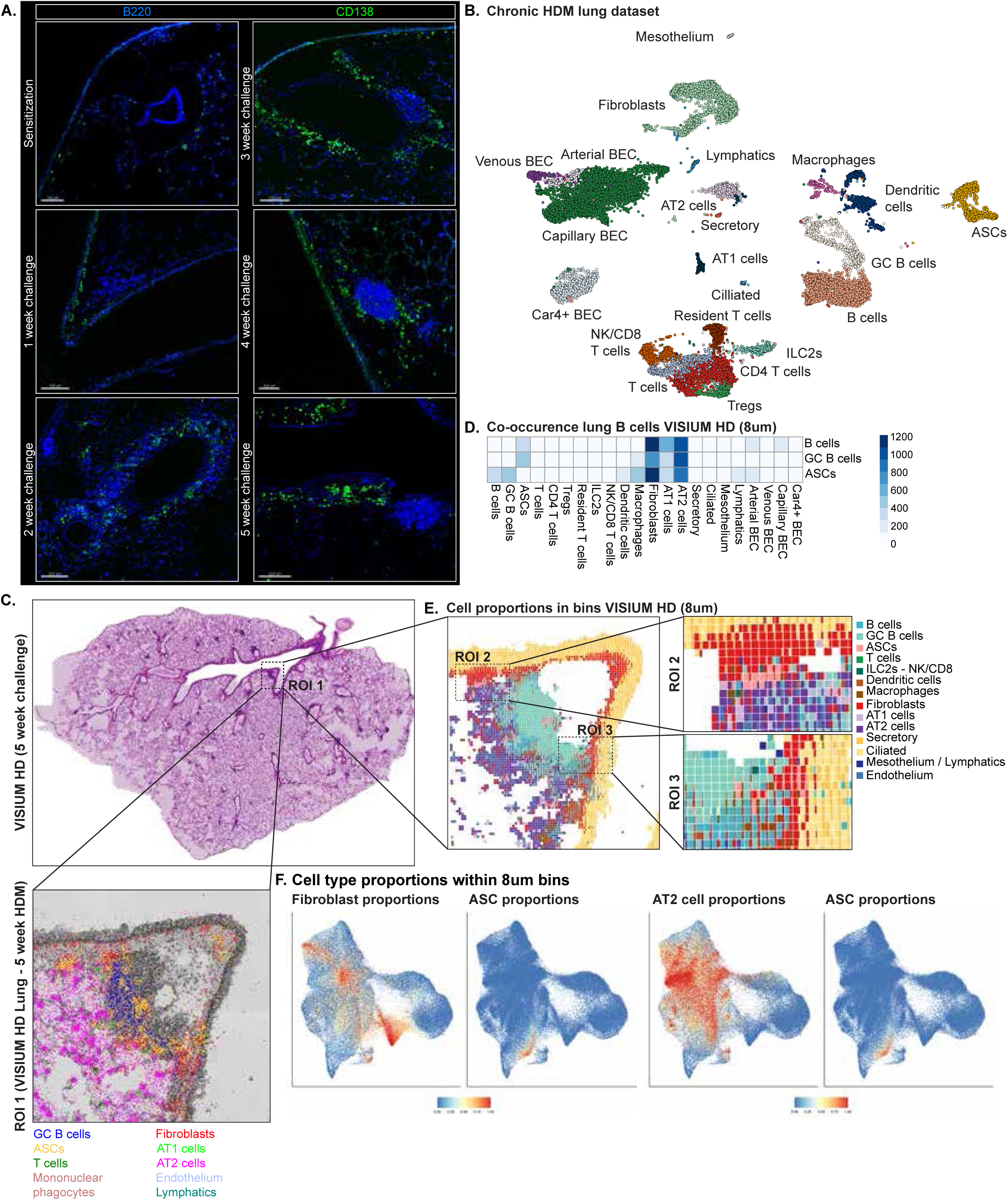
Lung ASCs exit TLO to find residence in the para-bronchial cuff, where they are in close proximity of lung fibroblasts and AT2 cells. A. Confocal imaging of ASC kinetics upon chronic HDM exposure. Scale bars: 100μm (sensitization, 4 weeks), 150μm (1 week) and 200μm (2 weeks, 3 weeks and 5 weeks) B. To F Single cell RNA sequencing and VISIUM HD on lungs exposed to HDM for 5 weeks. B. shows annotated UMAP of the chronic HDM lung dataset C. shows H&E image of the sequenced lung section with annotation plotted on ROI1. For spatial plotting simplicity, T cell clusters, endothelial clusters and macrophages and DCs from B, are combined D. Co-occurrence analysis of 8μm bins showing co-occurrence of B cell subsets to all other cell subsets and reveals close proximity of fibroblasts and AT2 cells with B cell subsets E. Cell proportions on 8μm bins on ROI1, with zoom on ROI2 and ROI3 that reveal co-occurrence of fibroblasts and AT2 cells with ASCs F. UMAPs showing cell type proportions within 8um bins: co-occurrence of fibroblasts and ASCs, and AT2 cells and ASCs

These observations suggest that persistent antigenic stimulation remodels the para-bronchial cuff into a niche that sustains lung ASCs. To identify the supportive cell types within this environment, we combined single-cell RNA sequencing (scRNA-seq) and VISIUM HD spatial transcriptomics of lungs after 5 weeks of HDM exposure. Sorted stromal cells and hashed immune cells from individual mice were loaded at a 1:1 ratio to ensure balanced representation (fig. S4, A and B). We generated a comprehensive single-cell dataset of lung populations under chronic HDM conditions and used it to deconvolute and estimate cell type composition of the 8μm bins of the VISIUM HD spatial dataset (Fig. 3, B and C). For visualization, related cell types were grouped (e.g., “T cells” include CD4⁺, regulatory, and tissue-resident subsets). Based on canonical marker expression, spatial mapping revealed distinct islands of TLO and ASCs embedded within alveolar type 2 (AT2)–rich regions (Fig. 3C; fig. S4, C–E). To assess potential niche interactions, we analyzed local co-occurrence of different cell types with the 8μm bins. After excluding intra-cluster overlaps, fibroblasts were most closely associated with AT2 cells (*48*), which in turn neighbored multiple endothelial subsets (fig. S4F). When focusing on TLO- and ASC-associated regions, we observed consistent co-localization of ASCs with fibroblasts and AT2 cells (Fig. 3D). High-resolution spatial mapping and cell proportion analyses confirmed these close associations (Fig. 3, E and F). Together, these results indicate that during chronic airway inflammation, ASCs relocate from TLO to para-bronchial cuff niches, where proximity to fibroblasts, and possibly AT2 cells, provides a supportive microenvironment for their maintenance and survival.

### Fibroblasts provide key survival cues to lung ASCs

To elucidate the signaling pathways supporting ASC responses within the para-bronchial cuff, we performed CellChat analysis on the chronic HDM lung dataset (Fig. 3B) (*49*). In line with spatial proximity data (Fig. 3), CellChat predicted strong interactions between fibroblasts or AT2 cells and B cell populations—including GC B cells and ASCs. Notably, fibroblast–ASC interactions were more frequent than those between AT2 cells and ASCs (fig. S5A). Among the predicted pathways, we identified significant enrichment of known ASC survival networks, including BAFF (*Tnfsf13b*), CXCL12, and IL-6 signaling (fig. S5B). Closer examination revealed that BAFF likely signals through TACI (*Tnfrsf13b*) on lung ASCs, as predicted contributions via BAFF-R (*Tnfrsf13c*) and BCMA (*Tnfrsf17*) were comparatively lower (Fig. 4A). Fibroblast-derived CXCL12 was predicted to act predominantly through CXCR4 on ASCs, since *Ackr3* was not expressed (Fig. 4B; fig. S5C). Fibroblast-derived IL-6 was predicted to signal via IL-6 trans-signaling through IL-6st, a mechanism previously implicated in lung transplant rejection (*50*) (Fig. 4C). Supporting this, VISIUM HD data showed that fibroblasts expressing *Tnfsf13b* (BAFF) or *Cxcl12* were preferentally located near ASCs, whereas *Il6* expression was detected only at low levels (Fig. 4D). Mesothelial cells, macrophages, and AT2 cells (in descending order of significance) were major predicted sources of APRIL (A proliferation-inducing ligand) directed toward ASCs (fig. S5D).

**Figure 4:**
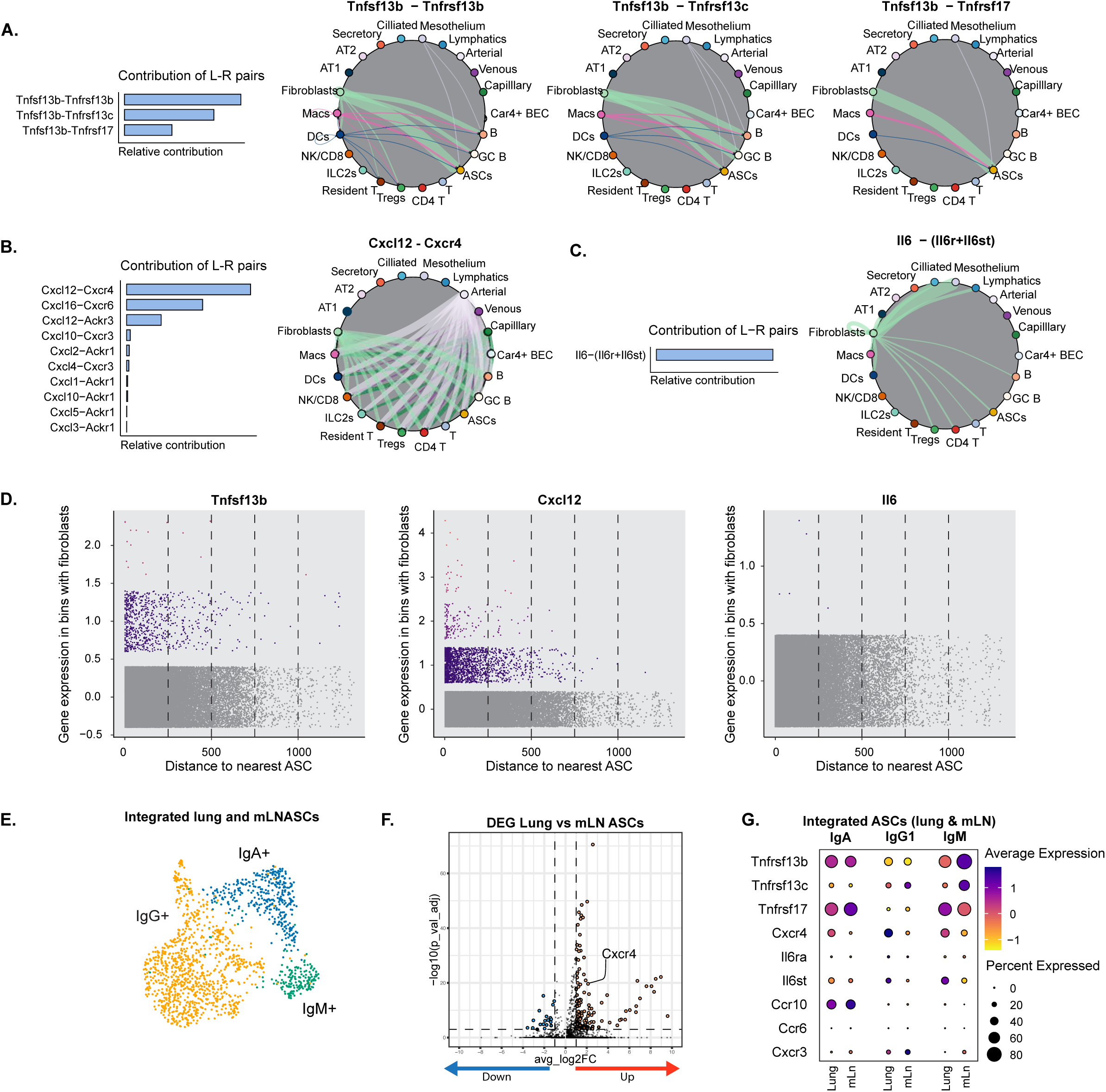
CellChat reveals lung fibroblast-ASC interactions enable ASC longevity and retention. A. To C CellChat analysis on the chronic HDM lung dataset shows strong interactions between fibroblasts and ASCs. A. TACI (*Tnfrsf13b*) mainly contributes to fibroblast-derived BAFF signaling to ASCs B. Predicted *Cxcl12* signaling from lung fibroblasts via *Cxcr4* on ASCs C. Fibroblast-derived *Il6* likely signals to ASCs via *Il6st* D. Gene expression (raw counts) of ASC supporting factors in bins with fibroblasts, correlated to distance to nearest ASC E. UMAP of integrated lung and mLN ASCs following chronic HDM exposure F. VolcanoPlot depicting differentially expressed genes (DEG) between lung and mLN ASCs, showing upregulation of *Cxcr4* on lung ASCs G. DotPlot showing expression of ASC survival and homing factors on lung and mLN ASCs per isotype

To compare receptor expression across mucosal and lymph node ASCs, we subsetted ASCs from lung and mLN within the scRNA-seq dataset (Fig. 4E). Differential expression analysis identified 128 genes upregulated and 19 downregulated in lung ASCs compared to mLN ASCs (log₂FC ≥ 1, adjusted pvalue ≤ 0.001), including the chemokine receptor *Cxcr4* (Fig. 4F). Indeed, *Cxcr4* was upregulated across all lung isotypes, whereas *Tnfrsf13b* (TACI) and *Tnfrsf17* (BCMA) were enriched in IgA⁺ and IgM⁺ ASCs, suggesting a role for these receptors in mucosal antibody responses (Fig. 4G). Consistent with prior reports (*1, 23, 51, 52*), *Ccr10* expression was restricted to IgA⁺ ASCs in both lung and mLN (Fig. 4G). Together, these findings indicate that fibroblasts in close proximity to ASCs provide a niche rich in BAFF, CXCL12, and IL-6, supporting the establishment and persistence of mucosal ASCs in the lung.

### IL-1 signaling instructs fibroblasts to support mucosal ASC residence

We next explored which upstream cues instruct fibroblasts to sustain ASC residency over time. CellChat analysis predicted IL-1β–IL-1R1 signaling interactions between macrophages, dendritic cells (DCs), and fibroblasts (Fig. 5A; fig. S6A). Specifically, conventional type 2 DCs (cDC2s), migratory cDC2s, classical monocytes, and interstitial macrophages (IMs) expressed high levels of *Il1b* (Fig. 5, B and C; fig. S6B). While our group previously demonstrated that IL-1 signaling in radioresistant stromal cells drives viral-induced TLO formation (*16*), its role in ASC organization and maintenance remains unknown.

**Figure 5:**
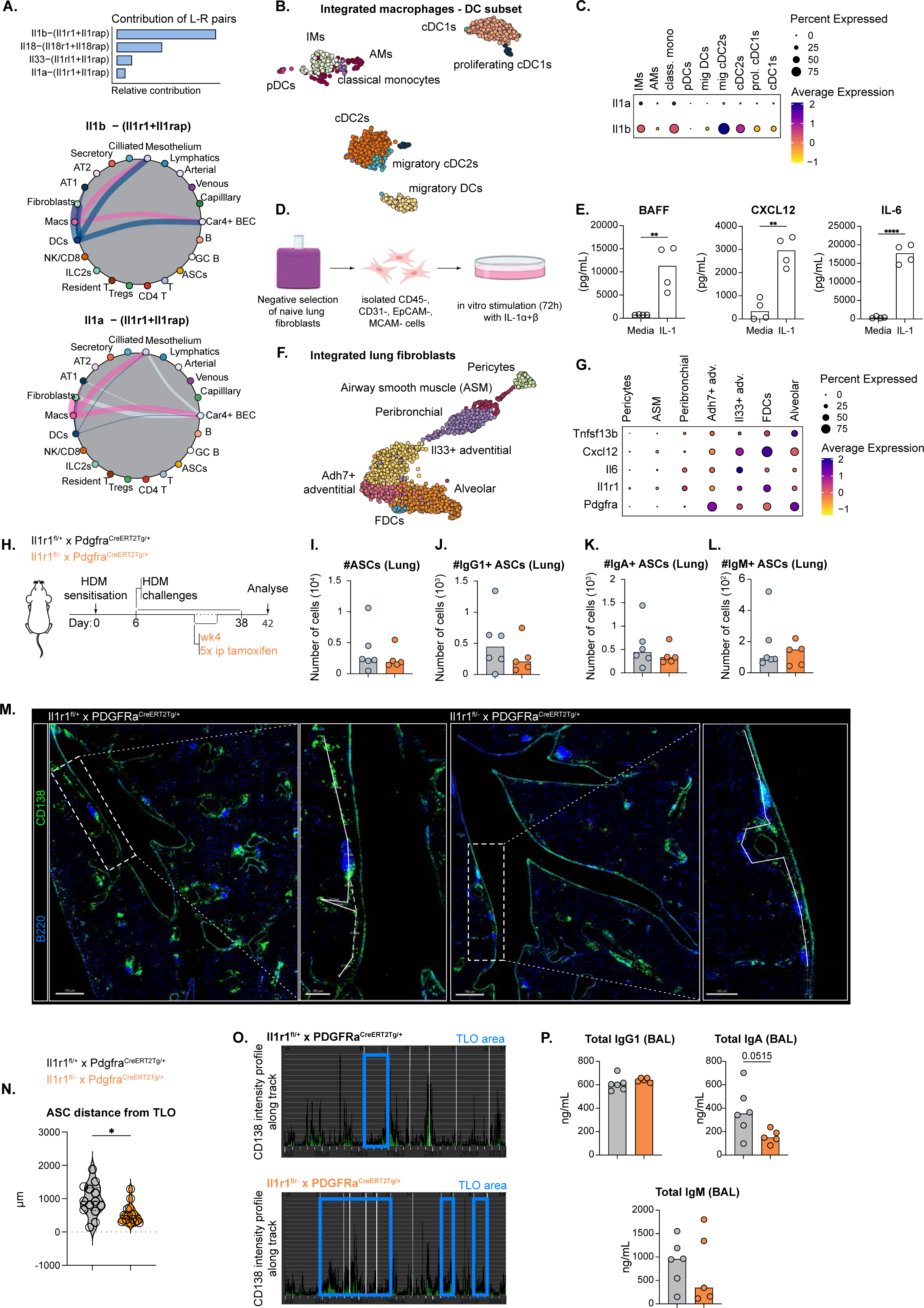
Fibroblast IL-1 signaling is required for ASC localization to para-bronchial niches in vivo. A. CellChat analysis on the chronic HDM lung dataset shows strong IL-1 interactions between macrophages/DCs to fibroblasts. B. UMAP of integrated lung macrophages and DCs following 5 weeks of HDM challenges C. DotPlot showing expression of *Il1a* and *Il1b* expression on lung macrophage and DC subsets D. Schematic of in vitro stimulation of isolated lung fibroblasts with recombinant IL-1 E. BAFF, CXCL12 and IL-6 secretion by IL-1 stimulated lung fibroblasts F. UMAP of integrated lung fibroblasts following 5 weeks of HDM challenges G. DotPlot showing expression of ASC survival factors *Tnfsf13b*, *Cxcl12* and *Il6*, and *Il1r1* and *Pdgfra* expression on lung fibroblast subsets H. Experimental set-up for temporal deletion of IL-1 signaling in Pdgfra+ fibroblasts during the chronic HDM protocol I. To L. Quantification of lung ASCs in Il1r1^fl/+^x Pdgfra^CreERT2Tg/+^ and Il1r1^fl/−^x Pdgfra^CreERT2Tg/+^ mice. I. showing total lung ASCs J. Quantification of IgG1^+^ lung ASCs K. Quantification of IgA^+^ lung ASCs L. Quantification of IgM^+^ lung ASCs M. Confocal analysis of ASC positioning into the para-bronchial cuff in Il1r1^fl/+^x Pdgfra^CreERT2Tg/+^ and Il1r1^fl/−^x Pdgfra^CreERT2Tg/+^ mice (scale bars: 700μm for overview images and 300μm for ROIs). ROIs show example of ASC track from TLO to calculate ASC-TLO distance and CD138 intensity profile. The dotted line on WT ROI (Il1r1^fl/+^x Pdgfra^CreERT2Tg/+^) was used for distance measurement. N. Quantification of ASC distance from TLO in Il1r1^fl/+^x Pdgfra^CreERT2Tg/+^ and Il1r1^fl/−^x Pdgfra^CreERT2Tg/+^ mice O. CD138 intensity profile along ASC tracks from TLO in Il1r1^fl/+^x Pdgfra^CreERT2Tg/+^ and Il1r1^fl/−^x Pdgfra^CreERT2Tg/+^ mice, showing that ASCs tend to reside more within the TLO in the absence of IL-1R signaling on Pdgfra+ fibroblasts. Blue boxes indicate TLO areas along the ASC track. ROIs for the tracks are indicated in M P. Total IgG1, IgA and IgM BAL levels in Il1r1^fl/+^x Pdgfra^CreERT2Tg/+^ and Il1r1^fl/−^x Pdgfra^CreERT2Tg/+^ mice For E, I-L, N, P, bars represent median with each symbol representing 1 replicate. P values are for Mann-Whitney U test (*P<0,05, **P<0,01, ***P<0,001). Data are representative of 3 independent experiments.

To test this, we isolated naive lung fibroblasts (*34*) and stimulated them with recombinant IL-1α and IL-1β (Fig. 5D). Combined IL-1α/β stimulation robustly induced secretion of BAFF, CXCL12, and IL-6 (Fig. 5E). Given the low expression of *Il1a* in our dataset (Fig. 5C), IL-1β likely dominates this response. Subclustering of fibroblasts in our scRNA-seq dataset confirmed substantial heterogeneity (*33–35*) (Fig. 5F; fig. S6C). ASC-supporting factors—*Tnfsf13b*, *Cxcl12*, and *Il6*—were highly expressed by FDCs within TLO and are likely to initiate ASC formation, as well as by IL33⁺ and *Adh7⁺* adventitial fibroblasts and alveolar fibroblasts (Fig. 5G). Expression of these factors closely correlated with *Il1r1* expression across these subsets, implicating IL-1 responsiveness as a key determinant of fibroblast function (Fig. 5G). Spatial analysis further revealed that macrophages and DCs expressing *Il1a/b* preferentially localized within 100 μm of fibroblasts (fig. S6D), supporting local IL-1–mediated fibroblast instruction. Collectively, these findings suggest that IL-1–responsive fibroblast subsets are instructed to orchestrate mucosal B cell responses and sustain ASC niches.

### Fibroblast IL-1 signaling is required for ASC positioning to para-bronchial niches in vivo

To functionally test the role of IL-1 signaling in fibroblast instruction, we crossed *Il1r1^flox^* mice with the *Pdgfra^CreERT2^* driver, which targets adventitial and alveolar fibroblasts (*33, 34, 46*) while enabling temporal gene deletion (Fig. 5H). To enhance deletion efficiency (*53*), *Il1r1^fl/fl^* × *Pdgfra^CreERT2^*mice were further crossed to *Il1r1^−/−^* mice, generating *Il1r1^fl/−^* × *Pdgfra^CreERT2^* littermates. Consistent with prior work (*46*), *Pdgfra^CreERT2^* efficiently targeted both fibroblast populations (fig. S6, E-G). Because IL-1 signaling is known to promote TLO formation (*16, 54*), we induced *Il1r1* deletion only after TLO GCs had formed to specifically assess effects on ASC maintenance (Fig. 2C; Fig. 5H).

Conditional *Il1r1* deletion in Pdgfra+ fibroblasts from week 4 onward did not affect TLO formation (fig. S6H). Total lung ASC numbers remained unchanged (Fig. 5, I-L; fig. S6,I and J); however, ASCs failed to migrate into para-bronchial cuffs, remaining confined to adventitial cuff TLO regions (Fig. 5, M-O, fig. S6K). This mislocalization correlated with a trend toward reduced BAL IgA and IgM, while IgG remained unaffected (Fig. 5, P). We find epithelial AT2 cells and secretory cells to a higher extent to express *Pigr* facilitating both IgA and IgM transcytosis into the airway lumen (fig. S6K) (*55*), we propose that impaired ASC migration to subepithelial para-bronchial niches limits mucosal IgA and IgM secretion. Although ASC positioning was affected, no changes in HDM-induced inflammation were observed (fig. S6L). Together, these data demonstrate that IL1-dependent fibroblast instruction is essential for ASC organization and positioning within para-bronchial niches enabling efficient antibody secretion into the airway mucosa. Thus, upon chronic inflammation, alveolar and adventitial fibroblasts orchestrate ASC residence and mucosal antibody airway immunity through IL1-dependent instruction.

## DISCUSSION

The airway mucosa represents a primary interface with the external environment and is constantly exposed to inhaled antigens, making it a critical site for immune surveillance and regulation (*56*). In chronic airway diseases, persistent antigen exposure drives prolonged inflammation, which can compromise lung function and gas exchange. Maintaining mucosal integrity is therefore essential. Using a model of continued inflammation induced by HDM extract, our findings reveal that prolonged allergen exposure induces bona fide germinal centers (GCs) within lung tertiary lymphoid organs (TLO), which generate mucosal antibody-secreting cells (ASCs). These ASCs preferentially localize to para-bronchial cuff niches adjacent to TLO, where prolonged inflammation instructs local fibroblasts to support ASC residence in an IL-1–dependent manner. This spatial positioning likely aligns ASCs with pIgR-expressing epithelial cells to facilitate targeted IgA and IgM secretion into the airway lumen.

Previous studies have implicated TLO as critical hubs for mucosal immunity, particularly in mice lacking conventional secondary lymphoid organs (SLOs) (*38, 43, 57*) or after CD11c-DTR–mediated TLO ablation (*15, 58*). Our single-cell RNA and BCR sequencing analyses indeed confirm that lung TLO GCs actively generate mucosal ASCs (*1, 23, 59*). Although clonal overlap between lung GC and ASC pools was limited in this polyclonal setting, these results, in line with recent publications (*18, 39*), provide direct evidence that mucosal ASCs can arise de novo from lung TLO GCs, clarifying the origin of airway antibodies in chronic allergic inflammation.

The concept of tissue niches orchestrating local immune responses has gained increasing attention (*19, 46, 50, 60*). Fibroblasts within these niches require reprogramming to accommodate tissue immunity (*33, 34*). In the lung, mucosal ASC homing has been mapped to influenza-infected alveoli, submucosal glands, and bronchovascular bundles, with CCR6, CXCR3, and CCR10 serving as key regulators (*1, 2, 17, 21, 22, 40, 51*). Consistent with these findings, CCR10 is highly expressed on IgA⁺ ASCs in our dataset. CXCL12 secreted by Gli1⁺Foxf1⁺Itga8⁻ fibroblasts near broncho-vascular bundles sustains mucosal ASCs, likely via IL-6 trans-signaling, in a chronic lung allorejection setting (*50*). We extend these observations by showing that, upon prolonged allergen exposure, IL-1 instructs phenotypically distinct fibroblast subsets to secrete CXCL12, thereby enabling ASC positioning into para-bronchial cuff niches.

Beyond homing, ASCs require docking and retention signals for their long term maintenance (*31*). Tissue-specific survival pathways vary: in bone marrow, BAFF and APRIL are essential, whereas in the lung, IL-6 and APRIL contribute (*40, 61, 62*). In agreement with prior reports, TACI predominantly supports mucosal ASC survival via BAFF rather than APRIL, which is minimally expressed in our dataset (*61, 62*). IgA- and IgM-secreting ASCs express BCMA (*Tnfrsf17*), implicating this receptor in their retention at mucosal sites. Although *Il6ra* was not detected transcriptionally on lung ASCs, IL-6 trans-signaling via *Il6st* may mediate ASC longevity. Additionally, tumors have been described to accumulate ASCs along fibroblast tracks(*63*). Collectively, these findings reveal a highly coordinated stromal–immune circuit in which IL-1–instructed fibroblast niches in the para-bronchial cuff guide ASC positioning, retention, and potentially IgA and IgM secretion. It will be interesting to elucidate if such IL-1 driven instruction also occurs in different sites of ASC residence, such as the nasal turbinates, lacrimal glands and the gut (*23*).

In the gut, mucosal IgA protects the epithelium against invading pathogens (*64, 65*), and similar protective roles have been described in the airways, olfactory mucosa, and brain (*1, 51, 52*). Additionally, IgA-deficient patients are at increased risk of allergic airway disease, and successful allergen immunotherapy correlates with elevated allergen-specific IgA in airway secretions (*66, 67*). Nevertheless, whether mucosal IgA directly exerts immunoregulatory functions in allergic airway disease remains debated (*68–74*). Although we find the mucosa to be enriched for IgA following continued HDM exposure while allergic inflammation conversely declined, an immunoregulatory role for mucosal IgA could not be confirmed. Attempts to demonstrate the role of IgA directly in IgA-deficient mice were confounded by compensatory increases in other isotypes, highlighting the need for improved IgA-specific models. However, the positioning of ASCs within the para-bronchial cuff might facilitate rapid antibody secretion within the airways, potentially enabling defence against new invading pathogens.

Taken together, our study describes how ectopic lymphoid organ formation and instruction of para-bronchial fibroblast niches promote local antibody production as an adaptive response to prolonged allergen exposure. Multiple fibroblast subsets contribute to ASC organisation and positioning in the para-bronchial cuff, highlighting the complexity and redundancy of this niche. Given the long-lived nature of fibroblasts, their altered functional state may have profound consequences for the persistence of proinflammatory and immunoregulatory immune cells, influencing chronic allergic disease. Whether these processes are dysregulated in allergic patients remains an important question for future investigation.

### Take-home message

Our study uncovers a mechanistic link between lung TLO GCs, para-bronchial fibroblast niches, and mucosal antibody production. IL-1–instructed fibroblasts position ASCs in subepithelial niches, enabling antibody secretion with a mucosal enrichment of IgA. These findings define a local stromal–immune circuit integrating continued inflammation with mucosal antibody responses, highlighting fibroblast-mediated niche remodeling as a critical regulator of lung immunity.

## MATERIALS AND METHODS

### Mice

C57BL/6 wild-type mice, aged 6-8 weeks, were purchased from Janvier (France). AID^CreERT2^xR26^TdTomato^ (strain #033897) reporter mice were a kind gift from Claude-Agnès Reynaud and Jean Claude Weill, Institut Necker Enfants Malades, France (*45*). Il1r^fl/fl^ (strain #028398) mice were obtained from the Jackson Laboratory. IL1r1^−/−^ mice (strain #003245) were crossed to PdgfraCreERT2^Tg/+^ mice, and Il1r1^−/+^ x Pdgfra^CreERT2Tg/+^ (strain #032770) mice were crossed to Il1r1^fl/fl^ (strain #028398, purchased from Jackson Laboratory) to obtain Il1r1^fl/−^x Pdgfra^CreERT2Tg/+^ mice.

Animals were housed in specific-pathogen free environment in the Laboratory Animal House at the VIB-UGent Center for Inflammation Research Center. All performed experiments were approved by the ethical committee of the VIB-UGent Center for Inflammation Research Center.

### Chronic house dust mite protocol and treatments

Mice, aged 6-8 weeks, were sensitized and challenged with HDM extract (ALK, Horsholm, Denmark). For HDM sensitisation on day 0, mice were anaesthetised using 2,5% isoflurane followed by intratracheal (i.t.) injection of 1μg HDM in 80μl of sterile Phosphate Buffered Saline (PBS). HDM challenges were performed starting 6 days after sensitisation, 3 times a week, with one day in between, for 5 weeks. For this, mice were again anaesthetised using 2,5% isoflurane, and 40μl of sterile PBS containing 10μg HDM was administered intranasally (i.n.). Mice were sacrificed 4 days after the last challenge via CO2 asphyxiation.

For AID enzyme labelling in AID^CreERT2^ x R26^TdTomato^ reporter mice, mice were i.p. injected with 2mg of Tamoxifen (Sigma, 75648-1G) in corn oil (Sigma, C8267-500ML) once, on day 27 of the chronic HDM protocol.

For LTβR-Fc signalling blockade, mice were intraperitoneally (i.p.) injected with 200μg LTβR-Fc on day 22, 29 and 36 of the chronic HDM protocol. Soluble LTβR-Fc was kindly provided by Carl F Ware (*75*) and was further produced in house by the VIB Protein Core under MTA with Sanford Burnham Prebys Research Institute, La Jolla, California, United States of America.

For temporal deletion of IL-1R on Pdgfra+ fibroblasts, Il1r1^fl/+^ x Pdgfra^CreERT2Tg/+^ and Il1r1^fl/−^ x Pdgfra^CreERT2Tg/+^ mice were i.p. injected with 2mg Tamoxifen in corn oil for 5 consecutive days, starting from day 27 until day 31 of the chronic HDM protocol.

### Organ collection and flow cytometric analysis

On the day of sacrifice, mice were intravenously (i.v.) injected with CD45 antibody (PE-eFluor610, clone 30-F11, Invitrogen, 61-0451-82) to allow blood from tissue discrimination. Following sacrifice, mice were terminally bled via inguinal vein cutting. Blood was collected, centrifuged and the isolated serum was stored at −20°C until further analysis. Broncho-alveolar lavage (BAL) fluid for ELISA analysis was collected by tracheal injection of 1ml BAL buffer (1L of 1x PBS + 1000ul of 0.5mEDTA), which was used to rinse the airways 3 times, followed by an additional injection of 2 times 1ml BAL buffer. BAL fluid was collected, centrifuged and supernatant was stored at −20°C until further analysis. BAL cell pellets were resuspended in the remaining 2ml BAL fluid, centrifuged and stained for flow cytometric analysis with the following antibodies in PBS: fixable viability dye eFluor506 (Invitrogen, 65-0866-18), CD11b V450 (clone M1/70, BD Biosciences, 560455), Ly-6G AF700 (clone 1A8, BioLegend, 127622), MHCII APC-eF780 (clone M5/114.15.2, eBioscience, 47-5321-82), Siglec-F PE (clone E50-2440, BD Biosciences, 552126), CD3e PE-Cy5 (clone 145-2c11, BioLegend, 100310), CD19 PE-Cy5 (clone eBio1D3(1D3), eBioscience, 15-0193-83), CD11c PE-Cy7 (clone N418, BioLegend, 117318).

mLN were isolated and collected in well plates containing RPMI. mLN were carefully pressed over a 100μm nylon mesh filter and transferred to a 96 u-well plate for staining. All lung lobes were harvested into RPMI. For the analysis of GC and ASCs, lungs were minced with scissors in FACS buffer (2.5g BSA + 1.14ml of 0.5M EDTA in 1L PBS), filtered over 100μm, centrifuged and red blood cells were lysed using osmotic lysis buffer before transferring to the staining plate. mLN and lung GC B cell surface staining in PBS was performed using the following antibodies: GL7 Pacific Blue (clone GL7, BioLegend, 144614), fixable viability dye eFluor506 (Invitrogen, 65-0866-18), IgM biotin (clone R6-60.2, BD Biosciences, 553406), CD138 BV711 (clone 281-2, BioLegend 142519), CD19 BV786 (clone 1D3, BD Biosciences, 563333), CD3e APC (clone 145-2C11, BioLegend, 100312), CD80 PE-Cy5 (clone 16-10A1, eBioscience, 15-0801-82), CD95 PE-Cy7 (clone Jo2, BD Biosciences, 557653), CD273 BUV379 (clone TY25, BD Biosciences, 565102). After incubating for 30 min at 4°C, cells were washed and stained with streptavidin BV605 (BD Biosciences, 563260) for another 15 minutes at 4°C. Next, cells were fixed and permeabilised (BD Cytofix/Cytoperm fixation 554722 and BD Perm/Wash Buffer 554723), and intracellularly stained in BD perm buffer with IgA Dylight488 (Abcam, ab97011) and IgG1 AF700 (clone RMG1-1, BioLegend, 406632). For detection of lung and mLN Tfh cells, the following antibodies were used: CD4 BV605 (clone RM4-5, BioLegend, 100548), CXCR5 BV650 (clone L138D7, BioLegend, 145517), PD-1 PE (clone J43, eBioscience, 12-9985-81) and Bcl6 AF647 (clone K112-91, BD Biosciences, 561525). For intracellular staining of the transcription factor Bcl6, cells were fixed and permeabilized using the eBioscience™ Foxp3 / Transcription Factor Staining Buffer Set (Fixation/Permeabilization Concentrate 00-5123; Fixation/Permeabilization Diluent 00-5223; Permeabilization Buffer (10X) 00-8333).

To study HDM-specific B cell responses and DarkZone/LightZone discrimination in the GC, lungs and mLN were minced in digestion buffer (20ug/ml Liberase (Sigma-Aldrich, 05 401 127 001), 0,001U/ul DNase 1 (Sigma-Aldrich 04 536 282 001) and 2%FCS in RPMI) and incubated at 37°C for 15 minutes while shaking. Cells were pipetted several times, and incubated for another 15 minutes at 37°C while shaking. Next, lungs were filtered over a 100μm nylon mesh filter and centrifuged. Red blood cell lysis was performed using osmotic lysis buffer. DarkZone and LightZone GC B cells were identified as described before (*6*) using CXCR4 AF647 (clone L276F12, BioLegend, 146504) and CD86 BV421 (clone GL1, BioLegend, 105032).

For identification and phenotyping of lung fibroblasts, lungs were inflated with Dispase digestion mix (5% FCS + 0,001U/ul DNase 1 (Sigma-Aldrich, 04 536 282 001) + 0.8-1mg/ml Dispase (Invitrogen, 17105-041) in RMPI). Tied off lungs were collected in wells filled with Dispase digestion mix and incubated at 37°C for 45 minutes. Next, lungs were minced in digestion buffer (20ug/ml Liberase (Sigma-Aldrich, 05 401 127 001), 0,001U/ul DNase 1 (Sigma-Aldrich 04 536 282 001) and 2%FCS in RPMI) and incubated at 37°C for 30 minutes while shaking. Lungs were resuspended several times before being transferred over a 100μm nylon mesh filter and centrifuged. Next, red blood cell lysis was performed using osmotic lysis buffer. Lungs were stained with the following antibodies to identify fibroblasts subsets: fixable viability dye APC-eF780 (Invitrogen, 65-0865-14), CD31 BV605 (clone 390, BD Biosciences, 740356), TER119 BV605 (clone TER-119, BioLegend, 116239), Epcam BV480 (clone G8.8, BD Biosciences, 746367), CD45 BV605 (clone 30-F11, BD Biosciences, 563053), Ly-51 FITC (clone 6C3, BioLegend, 108305), Sca1 Pacific Blue (clone D7, BioLegend, 108120), MHCII BV785 (clone MF/114.15.2, BioLegend, 107645), CD9 AF647 (clone MZ3, BioLegend, 124810), Pdpn PE-Cy7 (clone eBio8.1.1, Thermo Fisher Scientific, 25-5381-82), CD140a BUV737 (clone APA5, BD Biosciences, 741789). All samples were acquired on a BD LSRFortessa – 5 Laser system and the data was analysed with the FlowJo Software (v10.10.0).

### Single cell RNA-, CITE-R, TCR and BCR sequencing on lung and mLN cells

Adult C57BL/6 mice, aged 6-8 weeks, were subjected to the chronic HDM protocol as described above. For both single cell experiments sorting lung and mLN immune and stromal cells, 2 different digestion protocols were used: collagenase digestion (*76*) (RPMI + 2%FCS + 3mg/ml Collagenase IV (Sigma, C5138-100MG) and 40ug/ml DNase I (Sigma-Aldrich 04 536 282 001)) or dispase digestion (*77*). Following the two times 15 minute incubation at 37°C while shaking, tissue was transferred over a 100μm nylon mesh filter into FACS buffer. Single cell suspensions were centrifuged and red blood cell lysis was performed. A fraction of the lung and mLN cells were stained to check for adequate GC induction and stromal enrichment using the following antibodies: Pdpn AF488 (clone eBio8.1.1, eBioscience 53-5381-82), GL7 Pacific Blue (clone GL7, BioLegend, 144614), fixable viability dye eFluor506 (Invitrogen, 65-0866-18), CD31 biotin (clone 390, BD Biosciences, 740356), CD19 BV786 (clone 1D3, BD Biosciences, 563333), CD45 APC (clone 30-711, BioLegend, 103112), MHCII APC-eF780 (clone M5/114.15.2, eBioscience, 47-5321-82), CD95 PE-Cy7 (clone Jo2, BD Biosciences, 557653) and CD3e BUV737 (clone 145-2C11, BD Biosciences, 612771).

For FACS (Fluorescence-Activated Cell Sorting), CITE-seq and hashtag labelling, the remaining lung and mLN cells were counted, isolated and spun down. The cell pellet was resuspended and incubated for 30 min on ice with 50 µL of staining mix in PBS containing 0.04% BSA, the following antibodies for FACS (fixable viability dye eFluor506 (Invitrogen, 65-0866-18) and CD45 APC (30-F11, BioLegend, 103112), TruStain FcX Block (BioLegend, cat 101320), the mouse cell surface protein antibody panel containing 183 oligo-conjugated antibodies and 9 TotalSeq-C isotype controls (TotalSeq-C, BioLegend) (Supplementary Table6) and TotalSeq-C cell hashing antibodies (BioLegend) diluted 1:1000. Biological replicates (n = 6 for each organ) were multiplexed per lane using TotalSeq-C cell hashing antibodies. Sorted CD45^+^ and CD45^−^ single-cell suspensions were resuspended at an estimated final concentration of 200 - 1000 cells/µl and loaded on a Chromium GemCode Single Cell Instrument (10x Genomics) to generate single-cell gel beads-in-emulsion (GEM). The scRNA/Feature Barcoding/BCR/TCR libraries were prepared using the GemCode Single Cell 5’ Gel Bead and Library kit, version Next GEM 2 (10x Genomics) according to the manufacturer’s instructions. The cDNA content of pre-fragmentation and post-sample index PCR samples was analysed using the 2100 BioAnalyzer (Agilent). Sequencing libraries were loaded on an Illumina NovaSeq flow cell at VIB Nucleomics core with sequencing settings according to the recommendations of 10x Genomics (paired-end reads 28-10-10-90, 1% PhiX), pooled in a 75:10:10:5 ratio for the gene expression, TCR, BCR and antibody-derived libraries, respectively.

For the B cell lineage sequencing experiment, 6 adult C57BL/6 mice were subjected to the chronic HDM protocol as previously described. Lungs and mLN were harvested and processed (mincing for lung, smashing for mLN) as described above. Adequate induction of B cell responses in lung and mLN (GC B cells and ASCs) were checked in lung and mLN samples prior to FACS sorting, using the following antibodies: IgA Dylight488 (Abcam, ab97011), GL7 Pacific Blue (clone GL7, BioLegend, 144614), fixable viability dye eFluor506 (Invitrogen, 65-0866-18), CD138 BV711 (clone 281-2, BioLegend 142519), CD19 PE-Cy5 (clone eBio1D3(1D3), eBioscience, 15-0193-83), CD95 PE-Cy7 (clone Jo2, BD Biosciences, 557653).

Next, lung and mLN cells were counted, isolated and spun down for FACS and hashtag labelling. The cell pellets were resuspended and incubated for 30 min on ice with 50 µL of staining mix in PBS containing 0.04% BSA, containing the following antibodies for FACS (IgA Dylight488 (Abcam, ab97011), GL7 Pacific Blue (clone GL7, BioLegend, 144614), fixable viability dye eFluor506 (Invitrogen, 65-0866-18), CD138 BV711 (clone 281-2, BioLegend 142519), CD19 PE-Cy5 (clone eBio1D3(1D3), eBioscience, 15-0193-83), CD95 PE-Cy7 (clone Jo2, BD Biosciences, 557653), TruStain FcX Block (BioLegend, cat 101320) and TotalSeq-C cell hashing antibodies (BioLegend) diluted 1:1000. Biological replicates (n = 6 for each organ) were multiplexed per lane using TotalSeq-C cell hashing antibodies. Sorted GC B cells and ASCs from lung and mLN single-cell suspensions were resuspended at an estimated final concentration of 200 - 1000 cells/µl and loaded on a Chromium GemCode Single Cell Instrument (10x Genomics) to generate single-cell gel beads-in-emulsion (GEM). The scRNA/BCR libraries were prepared using the GemCode Single Cell 5’ Gel Bead and Library kit, version Next GEM 2 (10x Genomics) according to the manufacturer’s instructions. The cDNA content of pre-fragmentation and post-sample index PCR samples was analysed using the 2100 BioAnalyzer (Agilent). Sequencing libraries were loaded on an Illumina NovaSeq flow cell at VIB Nucleomics core with sequencing settings according to the recommendations of 10x Genomics (paired-end reads 28-10-10-90, 1% PhiX), pooled in a 75:10:10:5 ratio for the gene expression and BCR-derived libraries, respectively. FastQ files were mapped to the GRCm38.99 reference genome using CellRanger version 6.1.2 (10x Genomics). Preprocessing and processing of the data was done as described above.

### Sequencing data processing

FastQ files were mapped to the GRCm38.99 reference genome using CellRanger version 6.1.2 (10x Genomics). Preprocessing of the data was done by the scran and scater R package according to workflow proposed by the Marioni and Theis lab (*78, 79*). The subsamples labeled with hashing antibodies were identified using HTODemux and MultiSeqDemux by Seurat. Cells with less than 200 genes expressed and genes expressed in less than 3 cells were filtered out of the count matrix. Outliers were identified based on 3 metrics: number of expressed genes, library size and mitochondrial proportion. Cells 5 MADs (Median Absolute Deviation) away from the median value were filtered out for the number of expressed genes and library size, and 15 MADs were used as an upper limit for the mitochondrial proportion, resulting in lenient filtering. Detecting highly variable genes, scaling, finding clusters, and creating UMAP plots was done using the Seurat pipeline (see Rscripts for detailed description on the parameters used for each dataset).

Single-cell analysis was performed in R using Seurat v4.3.0(*80*). Three single cell experiments were performed and datasets were integrated according to the integration strategy (*81*). In 2 experiments, sorted stromal cells and hashed immune cells from individual mice chronically exposed to HDM were loaded for sequencing at a 50/50 ratio (Fig. 3B, fig. S 5A). To increase the number of clones for B cell lineage analysis, an additional experiment was set up where GC B cells and ASCs were sorted and hashed from 6 mice and loaded for RNA- and BCR-sequencing (fig. S 4N). In all experiments, tissue resident immune cells were isolated by exclusion of CD45^+^ IV labelled cells. Following integration of the 3 individual lung Seurat objects and unsupervised clustering, 21468 lung cells were retained (10392 lung cells from Lung sample1 and 11076 lung cells from Lung sample2). Sorted lung B cells were included in the integration yet left out for the analysis of the entire lung atlas to correct for B cell numbers. Cells from each dataset contributed to each cell cluster. Cell populations were identified using differentially expressed (DE) genes (fig. S 5B).

mLN datasets were integrated using an identical workflow (*81*). Immune cell clusters were identified based on elevated expression of canonical markers as described above (fig. S 5B). Although mLN samples were processed similarly as lung samples for scRNA-sequencing, in both stromal experiments, very few stromal cells could be retrieved from the mLN after loading the samples for sequencing, therefore the mLN data mainly consists of lymphocytes and very few fibroblasts.

For the B cell analysis, B cell clusters from the integrated lung and mLn datasets were integrated (*81*). scRNA-seq datasets from the B cell lineage experiment were included in the analysis. We retrieved 17831 B cells from 12 mice (5406 cells from 12 lung samples and 12425 cells from 12 mLN samples). B cell clusters were identified using representative markers (fig. S 4O). From this dataset, ASCs were subsetted (Fig. 4). In total, we retrieved 1565 ASCs (1153 lung ASCs from 12 samples and 412 mLN ASCs from 12 samples). ASCs clustered according to isotype expression (Fig. 4E) since Ig genes were taken into account for clustering.

For the BCR clonal analysis, BCRseq data from different samples was appended using rowbinding in R, followed by clonotype re-annotation by using the ImmCantation framework, following the method outlined in the instruction manual (*82*). Based on this information, a clonal analysis was performed by calculating the number of clonotypes shared between clusters. For this analysis, all samples were merged and clusters consisted of cells from the same tissue (lung or mLN), same cell type (LZ GC B cells, DZ GC B cells, ASC) and same antibody isotype.

Following lung macrophages and dendritic cell subsetting, 1203 cells were retained (671 cells from dataset lung sample1, and 532 cells from dataset lung sample2). Unsupervised clustering defined 9 clusters (fig. S 7B) (*83, 84*).

Following lung fibroblast subsetting, 1909 lung stromal cells were retained (827 stromal cells from dataset lung sample1 and 1082 from dataset lung sample2). Unsupervised clustering defined 8 clusters (fig. S 7D) (*33–35, 46, 85, 86*).

For visualisation purposes, we used the packages ggplot2 v3.4.0, ggrepel v0.9.2, scCustomize v1.1.1, viridis v0.6.2, and RColorBrewer v1.1.3. Number and percentage of cells per cluster were calculated using the package scCustomize v1.1.1. Venndiagrams were generated using the packages eulerr v7.0.0 and VennDiagram v1.7.3.

### Analysis of cell-cell communication using CellChat

Cell communication analysis was performed using the R package CellChat (*49*). We followed the RCTD vignette on the tool’s Github repository and ran all functions with default parameters, exept for the computation of communication probability (triMean of 0,05 was used, and min.cells was set at 10).

### Histology and confocal microscopy

Left lungs were inflated with a mixture of OCT and PBS (1:1) through tracheal injection, snap frozen in liquid nitrogen and stored at −80°C until further processing. Frozen sections were fixed with 2% PFA and aspecific binding was blocked through incubation with 0,1% mouse and rat serum in PBS. For TLO analysis, sections were stained using the following antibodies: GL7 Pacific Blue (clone GL7, BioLegend, 144614), CD35 BV480 (clone 8C12, BD Biosciences, 746338), Lyve-1 AF488 (clone ALY7, eBioscience, 53-0443-82), PNAd eFluor570 (clone meca-79, BioLegend, 120810), CD3e AF647 (clone 17A2, BD Biosciences, 557869) B220 AF700 (clone RA3-6B2, BD Biosciences, 557957). For ASC imaging, sections were stained overnight using primary labelled antibodies: GL7 Pacific Blue (clone GL7, BioLegend, 144614), B220 BV480 (clone RA3-6A2, BD Biosciences, 565631), IgA Dylight650 (abcam, ab97014), IgG1 AF700 (clone RMG1-1, BioLegend, 406632). IgM staining was performed using IgM biotin (clone R6-60.2, BD Biosciences, 553406) followed by streptavidin AF488 (eBioscience, S32354).

For quantification of ASC distance from TLO, measurement points were added in the Imaris analysis software, defining the ASC track delineating TLO exit and the intensity profile for the ASC marker CD138 was obtained from the software. All images were captured with the Leica Stellaris 8 microscope and analysed using the Imaris Software (v10.2).

### Collecting lung samples for VISIUM HD

C57BL/6 wild-type mice, aged 6-8 weeks, were purchased from Janvier (France). Following chronic HDM exposure, mice were euthanised and lungs were inflated with a pre-cooled mixture of OCT compound and PBS (1:1 ratio). Individual lung lobes were isolated, briefly rinsed with pre-cooled OCT before transferring into a cryomold filled with pre-cooled OCT. Cryomolds were frozen in isopentane, and stored at −80°C until further processing. Fresh frozen lung sections (10 µm thick) were cut and mounted onto Nexterion® Slides (Schott AG, Germany), following the guidelines provided in the 10x Genomics Visium HD Fresh Frozen Tissue Preparation Handbook (CG000763, Rev C). Slides were stored at −80 °C until subsequent processing. Slide processing began with fixation as described in the aforementioned protocol. RNA capture and library preparation were performed according to the Visium HD protocol (CG000685, Rev C). Sequencing was carried out with a read depth of 242 million reads (70.9% sequencing saturation). Data processing was performed using Space Ranger.

### Processing of VISIUM HD lung data

We applied the deconvolution tool RCTD (*87*), implemented in the spacexr package (v2.2.0), to estimate cell type compositions in the lung Visium HD slide at the 8 µm resolution. As reference for the deconvolution, we used raw scRNA-seq counts from the chronic HDM lung dataset which had been annotated into 22 clusters. We followed the RCTD vignette on the tool’s Github repository and ran all functions with the default parameters, including UMI_min=100, which indicates that a minimum of 100 UMIs is required for a bin to be retained in the analysis. This filtered out ∼72% of the bins, leaving ∼140,000 bins for the analysis. We used code optimizations implemented by 10x Genomics to speed up the computation (dmcable/spacexr#206). This optimization is available only for doublet_mode=’doublet’ where a maximum of two cell types can be assigned per bin. After obtaining the proportions, cell type co-occurrence was calculated by tallying the number of times pairs of cell types were predicted within the same bin. Therefore, bins classified as singlets were excluded, and the magnitude of the proportions were not taken into account in this calculation.

### In vitro culture of magnetically sorted lung fibroblasts

Naive C57BL/6 mice were euthanised and lungs were processed using the Dispase digestion method as described above. Lung samples were incubated with the following antibodies to allow magnetic negative selection as described previously(*34*): CD45 APC (30-F11, BioLegend, 103112), CD31 APC (MEC 13.3, BD Biosciences, 551262), EpCAM APC (G8.8, BioLegend, 118214), MCAM APC (ME-9F1, BioLegend, 134712). Magnetic negative selection using anti-APC MicroBeads (Miltenyi, 130-090-855) was performed. 2×10e5 cells were seeded into 96 well plates and cultured in DMEM with 10% FCS and 1% penicillin-streptomycin for 24 hours. After adherence of fibroblasts, any remaining red blood cells were washed away. Following medium starvation, medium was changed for stimulation with medium containing IL-1 mix (10ng/ml mouse IL-1a (R&D systems, 400-ML-025) and 10ng/ml mouse IL-1b (R&D systems, 401-ML-005)) for 72 hours. Cell cultures were performed under standard culture conditions (37°C, 5% CO_2_).

### ELISA

Total immunoglobulin (Ig) levels in BAL and serum were evaluated using the following antibodies: anti-IgG1 capture antibody (BD Biosciences, 553445), anti-IgG1 biotinylated detecting antibody (BD Biosciences, 553441), anti-IgA capture antibody (BD Biosciences, 556969), anti-IgA biotinylated detecting antibody (BD Biosciences, 556978), anti-IgM capture antibody (BD Biosciences, 553435), anti-IgM biotinylated detecting antibody (BD Biosciences, 553406), anti-IgE capture antibody (BD Biosciences, 553413), anti-IgE biotinylated detecting antibody (BD Biosciences, 553419). Standard curves were measured using mouse IgG1 standard (BD Biosciences, 557273), mouse IgA standard (BD Biosciences, 553476), mouse IgM standard (BD Biosciences, 553472) and mouse IgE standard (BD Biosciences, 557079). CXCL12, CXCL13 and BAFF concentrations in culture supernatant were evaluated with the Duoset elisa kits from R&D Systems. IL-6 concentrations in culture supernatans were evealuated using the Ready-SET-Go! ELISA kits (Thermo Fisher Scientific).

IL-4, IL-5, IL-10, IL-13 concentrations in BAL were evaluated using the Ready-SET-Go! ELISA kits (Thermo Fisher Scientific).

For all ELISA analyses, absorbance was read at 450nm and 650nm as reference with a Perkin Elmer multilabel counter and data were collected with Wallac 1420 Manager software (PerkinElmer).

### ELISpot Assays

C57BL/6 wild-type mice, aged 6-8 weeks, were purchased from Janvier (France). Following chronic HDM exposure, mice were intravenously injected with CD45 biotin antibody (clone 30-F11, Thermo Fisher Scientific, 13-0451-85). Lungs and mLN were harvested and processed as described above. ASCs were enriched using negative magnetic enrichment using the following antibodies: CD3e biotin (145-2C11, Thermo Fisher Scientific, 13-0031-85). ELISpot assays were performed according to following standard protocols: ELISpot Flex: Mouse IgG1 (ALP) (Mabtech, 3821-21), ELISpot Flex: Mouse IgA (ALP) (Mabtech, 3865-2A), ELISpot Flex: Mouse IgM (ALP) (Mabtech, 3885-2A). Briefly, ELISpot plates (Mabtech, 3654-WP-10) were coated with the following antibodies: anti-mouse IgG1 (Mabtech, MT24/JC5-1), anti-mouse IgA (Mabtech, MT45A), anti-mouse IgM (Mabtech, MT6A3) and incubated overnight at 4°C. Following isolation, 45 000 cells per well were incubated in Assay Medium (RPMI 1640 supplemented with 2nM L-glutamine, 10% FCS, 1% Penicillin (100ug/mL) +Streptomycin (100ug/mL), 8um HEPES, 50uM 2-mercaptoethanol) for 21h under standard culture conditions (37°C, 5% CO_2_). ASCs were detected using the following detection antibodies: Anti-IgG1 (Mabtech, MG1), anti-IgA (Mabtech, MT39A) and anti-IgM (Mabtech, MT9A2). Detection antibodies were revealed with streptavidin-alkaline phosphatase (Mabtech, 3310-10-1000) and developed with BCIP/NBT (Mabtech, 3650-10). ELISpot plates were imaged and counted using the Mabtech Iris plate reader (Mabtech).

### Statistical analysis

Kinetics data are represented as the mean of the data ± standard error of the mean (SEM) and statistical significance between groups and timepoints was calculated with 2-way-ANOVA using GraphPad Prism Software (v10.2.0 (355), GraphPad Software, La Jolla, Calif). For other analyses (and as indicated in the figure legend), data are represented as individual values with bars representing median. Statistical significance between groups was calculated with Mann-Whitney U test (when comparing 2 groups) or ordinary one-way ANOVA with Sidak’s multiple comparisons test (when comparing more than 2 groups) using GraphPad Prism software. Differences between groups were considered significant at P values of 0.05 or less, 0.01 or less, 0.001 or less, as indicated in the figure legend. To ensure sufficient statistical power, sample sizes were determined empirically using G*power (v3.1). Mice were randomly assigned to experimental groups. Experiments were not performed blinded. No data was excluded from the analysis.

### Textual optimization

ChatGPT was used for final textual optimization.

## List of Supplementary materials

### Supplementary Figures

Supplementary Fig. 1 to 6

### Auxiliary Supplementary Material files

Supplementary Table1_Chronic-HDM-lung-mLn-Bcells_markers.xlsx

Supplementary Table2_Chronic-HDM-lung-atlas_markers.xlsx

Supplementary Table3_Chronic-HDM-lung-mLn-ASCs_markers.xlsx

Supplementary Table4_Chronic-HDM-lung-Macrophages-DCs_markers.xlsx

Supplementary Table5_Chronic-HDM-lung-Fibroblasts_markers.xlsx

Supplementary Table6_TotalSeqC antibodies.xlsx

MDAR Reproducibility Checklist

Supplementary Tables1-5 include excel datasheets showing DE genes per annotated cluster for each dataset that is used in the manuscript. Supplementary Table6 shows TotalSeq antibodies used for CITE-sequencing.

## Acknowledgements

We thank the technicians of the Lambrecht-Hammad lab, VIB Flow Core, the VIB Bioimaging Core and the VIB Protein Core. We thank the VIB Single Cell Core for support regarding single-cell technologies (single-cell.be) and the VIB Spatial Catalyst core for the support regarding spatial transcriptomics technologies. We thank the VIB Nucleomics Core for support and access to the instrument park (vib.be/core-facilities).

We thank Carl F. Ware for providing the LTβR-Fc construct. We thank the Animal facility at VIB-UGent for their excellent animal care. We thank Jannes Gavel for carefully revising the manuscript. Lastly, we would like to thank all colleagues for the insightful discussions and feedback.

## Funding

Fonds voor Wetenschappelijk Onderzoek Vlaanderen (FWO) PhD Fellowship fundamental research grants: 11D9623N (IL), 3F015119 (JD)

Fonds voor Wetenschappelijk Onderzoek Vlaanderen (FWO) junior postdoctoral researcher grant 3E011619 (ASB)

Fonds voor Wetenschappelijk Onderzoek Vlaanderen (FWO) senior postdoctoral researcher grants: 3E002820 (SV)

Fonds voor Wetenschappelijk Onderzoek Vlaanderen (FWO) MSCA Seal of Excellence fellowship (312ZZR22) (US)

RESPIRE4 Marie Sklodowska-Curie Postdoctoral Research Fellowship R4202007-00839 (US) LEO Foundation Serendipity grant (EH)

Fonds voor Wetenschappelijk Onderzoek Vlaanderen (FWO) PhD Fellowship Strategic Basic research grant S006825N (KH)

Ghent University Special Research Fund grant BOF21-DOC-105 (CS)

RESPIRE4 Marie Sklodowska-Curie Postdoctoral Research Fellowship (APB)

Flanders AI Research Program grant: 174K02325 (RS)

Flanders AI Research Program grant: 174K02325, Ghent University Special Research Fund grant: 01G03524, and the Belgian Excellence of Science (EOS) program grant: 3G0I2722 (YS)

Fonds voor Wetenschappelijk Onderzoek Vlaanderen (FWO) Research project grants: 3G0A7422 (PG, SV), 3G062218W (HH)

iBOF-FWO grant 01IB0423 (BNL)

BOF Methusalem grant 01M01521 (BNL)

Bijzonder Onderzoeksfonds Universiteit Gent (BOF): bof/baf/2y/2024/01/017 (SV)

## Author contributions

Conceptualization: IL, PG, HH, BNL and SV

Methodology: IL, BNL and SV

Investigation: IL, JD, ASB, US, APB, BNL and SV

Visualization: IL, KH, CS, RS, RB and SV

Funding acquisition: IL, HH, PG, BNL and SV

Project administration: IL, BNL, and SV

Supervision: HH, BNL RS, YS, and SV

Writing – original draft: IL and SV

Writing – review & editing: IL, ASB, US, APB, RS, BNL and SV

## Data and material availability

LTβR-Fc was obtained under MTA with Dr. Carl F Ware (Sanford Burnham Prebys Research Institute, La Jolla, California, United States of America). All other data are available in the main text or the supplementary materials.

