## Supplementary figures for "IL-1 instructs para-bronchial cuff fibroblasts to organize lung antibody secreting cell niches during continued antigen exposure"

**Lammens et al. 2026**

**Supplemental figures**

Supplemental figure 1 - Related to figure 1

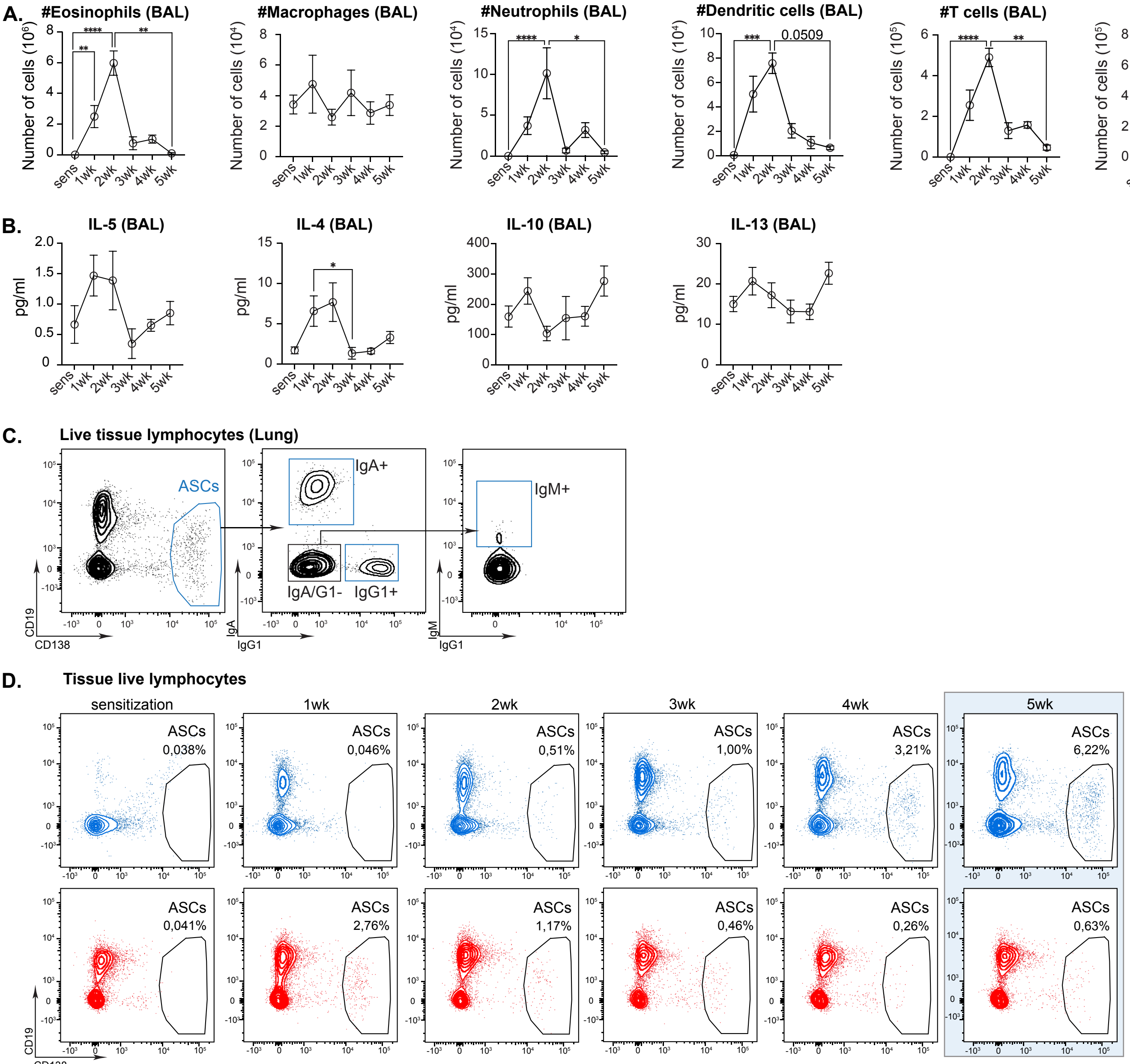

**Fig. S1: Upon chronic HDM exposure, lung ASCs are enriched in the airway mucosa while allergen-induced inflammation wanes over time.**

A. Kinetics analysis of allergen-induced airway inflammation showing BAL eosinophils, BAL macrophage, BAL neutrophil, BAL dendritic cell, BAL T cell and BAL B cell numbers
B. Kinetics analysis of type 2 cytokines in BAL
C. Gating strategy to identify ASCs
D. Flow cytometric kinetics of lung (blue) and mLN (red) ASCs during chronic HDM exposure

For A-B, data are shown as means + sem (n=4 for each timepoint). P values are for one-way-ANOVA (*P<0,05, **P<0,01, ***P<0,001). Data are representative of 3 independent experiments. Related to figure 1.

Supplemental figure 2 - Related to figure 2

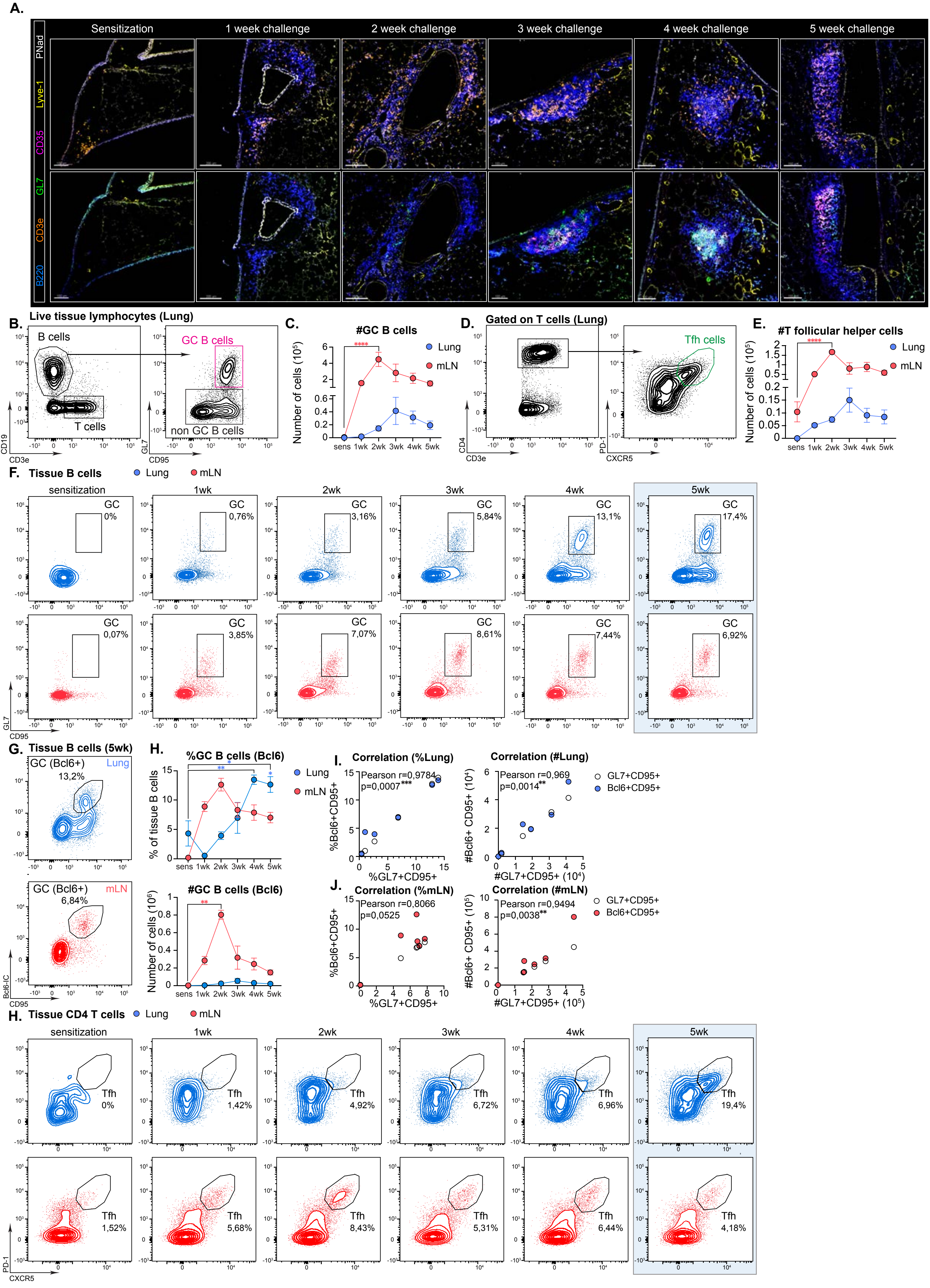

**Fig. S2: TLO develop within the adventitial cuff starting from 3 weeks of HDM challenges.**

A. Confocal imaging of TLO kinetics upon chronic HDM exposure showing the formation of B cell follicles (blue) with GC zones (green) and FDCs (pink) from 3 weeks of HDM exposure, T cells (orange), lymphatics (yellow) and HEVs (white). Scale bars: 100μm (sensitization, 3 weeks, 4 weeks), 150μm (5 weeks) and 200μm (1 week and 2 weeks)
B. Gating strategy to identify GC B cells in lung (5 week timepoint)
C. Quantification of GC B cell kinetics in lung (blue) and mLN (red)
D. Gating strategy to identify T follicular helper T cells (Tfh) in lung (5 week timepoint)
E. Quantification of Tfh cell kinetics in lung (blue) and mLN (red)
F. Flow cytometric kinetics of lung (blue) and mLN (red) GC B cells
G. Flow cytometric analysis of Bcl6 expression in GC B cells in lung (blue) and mLN (red), 5 week timepoint shown.
H. Quantification of G
I. To J. Correlation of GC B cells gated via GL7 vs Bcl6 during the chronic HDM protocol for lung (blue, H) and
J. mLN (red, J)
K. Flow cytometric kinetics of lung (blue) and mLN (red) Tfh cells

For C, E and H, data are show as means + sem (n=4 for each timepoint). P values are for 2-way-ANOVA (C, E and H) (*P<0,05, **P<0,01, ***P<0,001). Data are representative of 3 independent experiments. Related to figure 2.

Supplemental figure 3 - related to figure 2

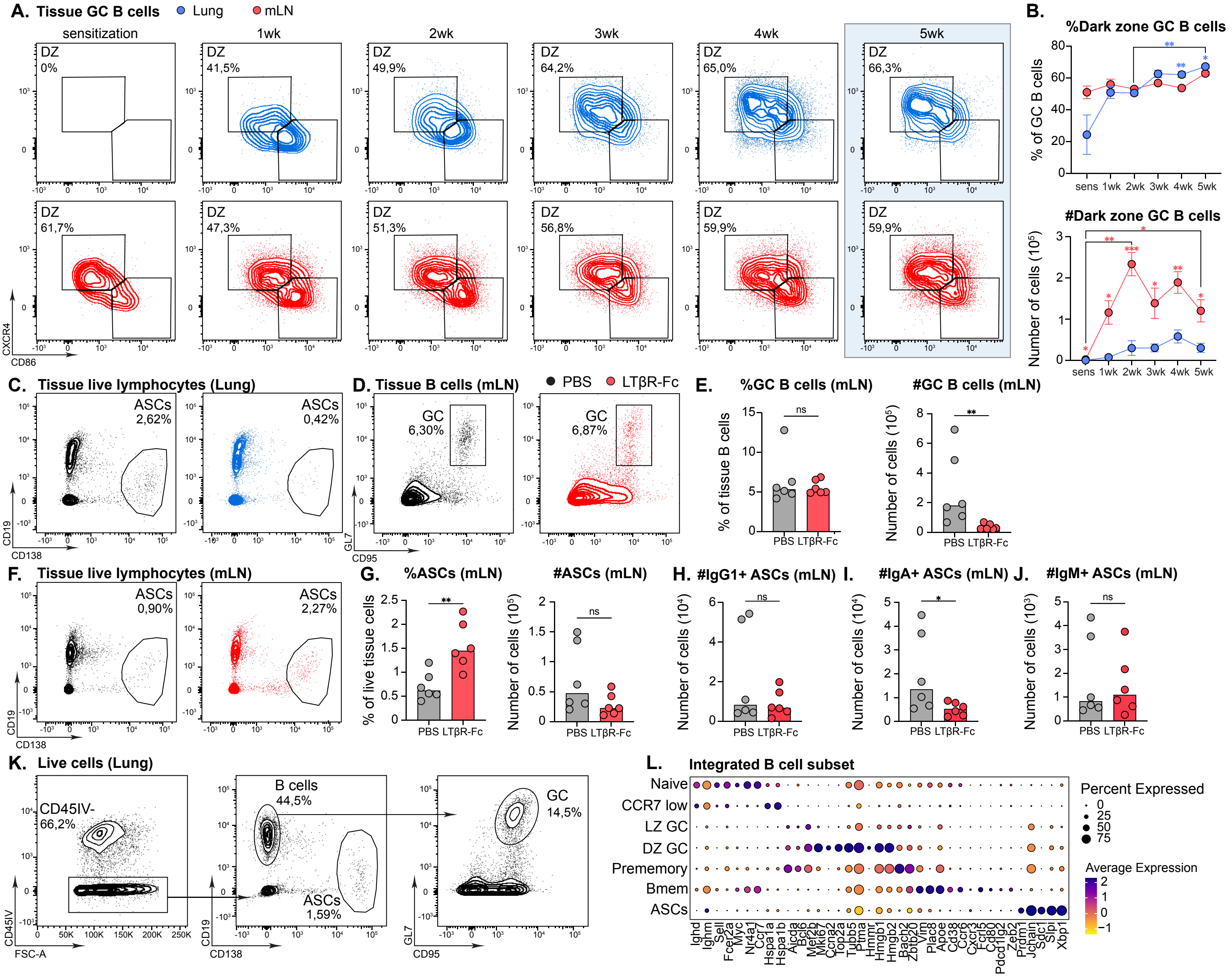

**Fig. S3: Chronic HDM exposure induces mature TLO GC in the lung that generate ASCs**

A. Flow cytometric kinetics of lung (blue) and mLN (red) dark zone and light zone GC B cells
B. Quantification of A
C. Flow cytometric analysis of lung ASCs following PBS vs LTβR-Fc administration
D. Flow cytometric analysis of mLN GC B cells upon PBS vs LTβR-Fc administration
E. Quantification of D
F. Flow cytometric analysis of mLN ASCs following PBS vs LTβR-Fc administration
G. To J Quantification of F
H. Quantification of mLN $IgG1^+$ ASCs following PBS vs LTβR-Fc administration
I. Quantification of mLN $IgA^+$ ASCs following PBS vs LTβR-Fc administration
J. Quantification of mLN $IgM^+$ ASCs following PBS vs LTβR-Fc administration
K. Gating strategy for lung and mLN B cell lineage sorting for BCRsequencing
L. DotPlot showing curated list of genes for B cell annotation

For B, data are show as means + sem (n=4 for each timepoint). For E and G-J, bars represent median with each symbol representing 1 replicate. P values are for 2-way-ANOVA (B) and Mann-Whitney U test (E and G-J) (*P<0,05, **P<0,01, ***P<0,001). Data are representative of 3 independent experiments. Related to figure 2.

Supplemental figure 4 - Related to figure 3

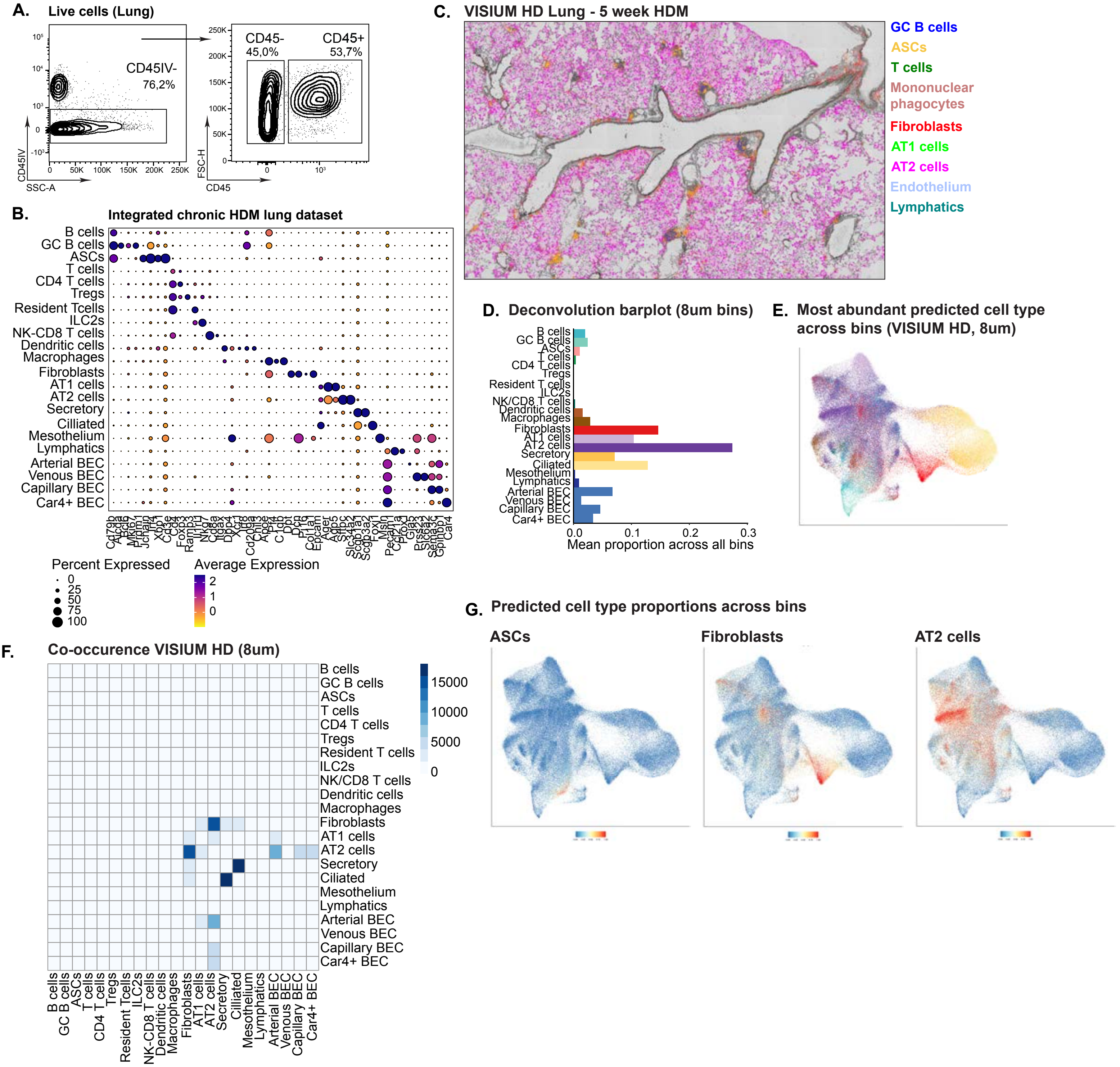

**Fig S4: Chronic HDM lung VISIUM HD data show close co-occurrence of lung ASCs to fibroblasts and AT2 cells**

A. Gating strategy for lung and mLN immune and stromal cell sorting for scRNAsequencing
B. DotPlot showing curated list of genes for chronic lung HDM dataset annotation
C. Annotation of cell clusters on chronic lung HDM VISIUM HD based on the chronic HDM lung dataset
D. Deconvolution barplot on 8μm bins showing mean proportions of cell types across VISIUM HD lung slide
E. Lung VISIUM HD UMAP on 8μm bins showing the most abundant cell type predicted in each bin
F. Lung VISIUM HD co-occurrence matrix on 8μm bins showing the number of times a cell pair is present in a bin. Only bins with >100 UMIs were included in the analysis. Co-occurrence of the same cell type in the bin was left out of the analysis
G. Lung VISIUM HD UMAP on 8μm bins showing the predicted cell type in each bin, for ASCs, fibroblasts and AT2 cells

Related to figure 3

Supplemental figure 5 - Related to figure 4

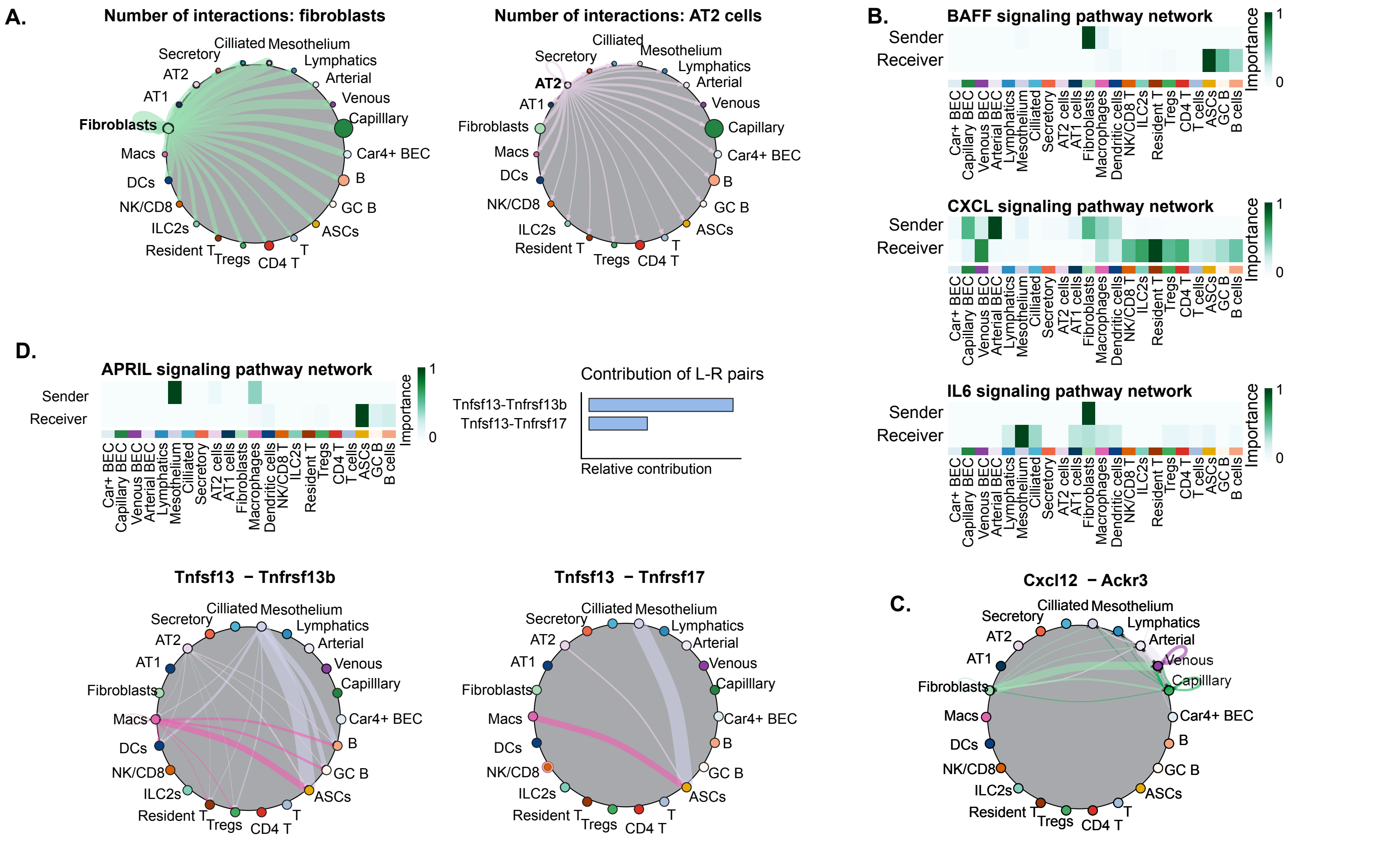

**Fig S5: CellChat reveals significant interactions between lung fibroblasts and ASCs, possibly enabling lung ASC survival.**

A. CellChat analysis showing number of interactions from fibroblasts and AT2 cells. Thickness of connecting lines corresponds to the number of interactions
B. Heatmap from CellChat analysis shows the relative importance of each cell type as sender and receiver for BAFF signaling, CXCL signaling and IL6 signaling
C. Predicted *Cxcl12-Ackr3* interactions mainly occur between fibroblasts and venous endothelium
D. CellChat analysis predicts significant interactions for the APRIL signaling network between mesothelium and macrophages (in order of importance), to ASCs, that mainly sense APRIL via TACI

Related to figure 4

### Supplemental figure 6 - Related to figure 5

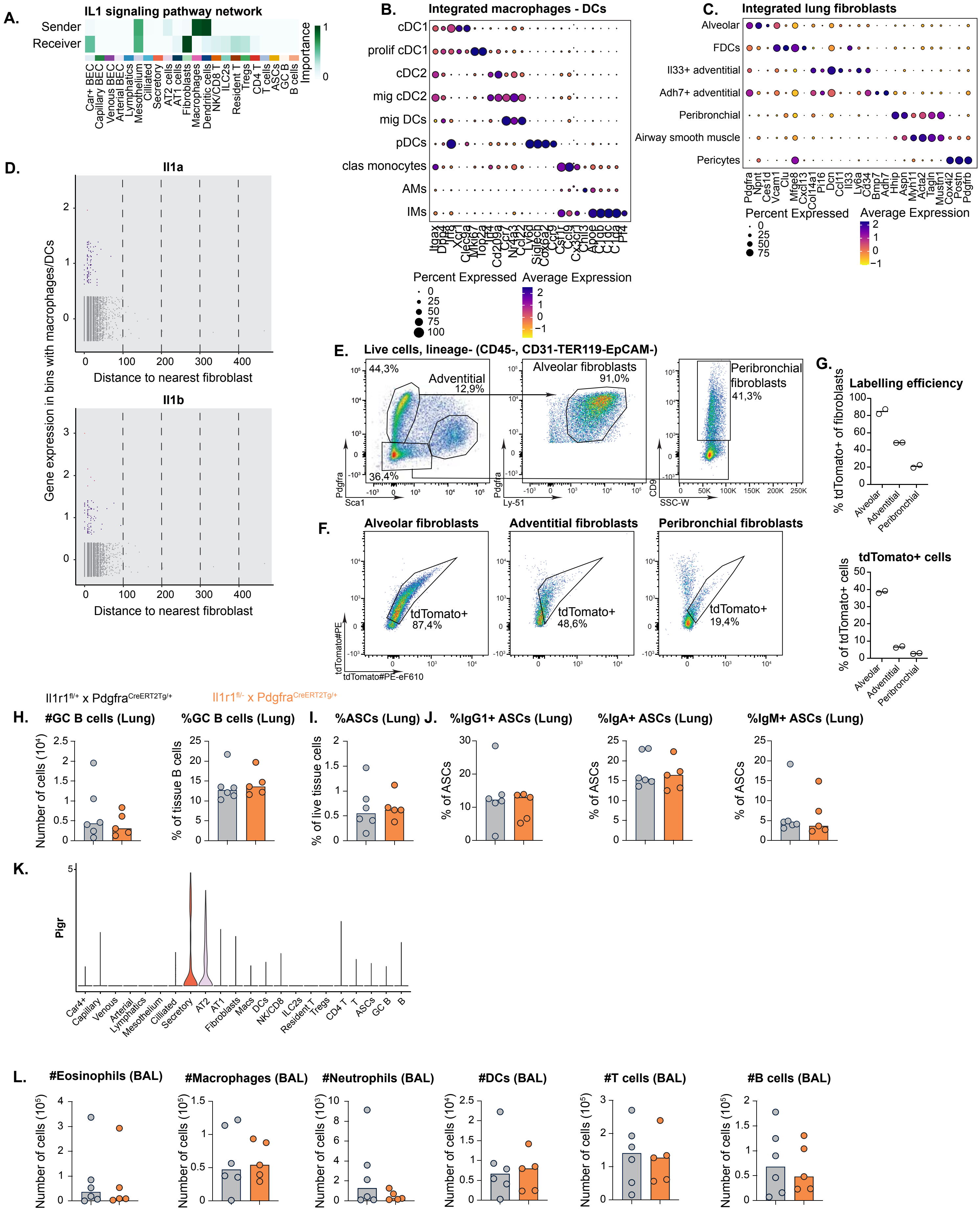

**Fig. S6: Fibroblast IL-1 signaling is required for ASC localization to para-bronchial niches in vivo.**

A. Heatmap from CellChat analysis shows the relative importance of each cell type as sender and receiver for IL1 signaling
B. DotPlot showing curated list of genes for annotation of integrated lung macrophages and DC subset
C. DotPlot showing curated list of genes for annotation of integrated lung fibroblast subset
D. Gene expression (raw counts) of *Il1a* and *Il1b* in bins with macrophages and DCs, correlated to distance to nearest fibroblast
E. Gating strategy to identify lung fibroblast subsets
F. Flow cytometric analysis of tdTomato expression in lung fibroblast subsets of Pdgfra$^{CreERT2}$ reporter mice.
G. Quantification of F
H. Quantification of lung GC B cells in Il1r1$^{fl/+}$x Pdgfra$^{CreERT2Tg/+}$ and Il1r1$^{fl/-}$x Pdgfra$^{CreERT2Tg/+}$ mice
I. Percentage of lung ASCs in Il1r1$^{fl/+}$x Pdgfra$^{CreERT2Tg/+}$ and Il1r1$^{fl/-}$x Pdgfra$^{CreERT2Tg/+}$ mice
J. Percentage of IgG1$^{+}$, IgA$^{+}$ and IgM$^{+}$ lung ASCs in Il1r1$^{fl/+}$x Pdgfra$^{CreERT2Tg/+}$ and Il1r1$^{fl/-}$x Pdgfra$^{CreERT2Tg/+}$ mice
K. VlnPlot showing highest *Pigr* expression in secretory cells followed by AT2 cells in the chronic HDM model
L. Quantification of allergen-induced airway inflammation in Il1r1$^{fl/+}$x Pdgfra$^{CreERT2Tg/+}$ and Il1r1$^{fl/-}$x Pdgfra$^{CreERT2Tg/+}$ mice, showing BAL eosinophils, macrophage, neutrophil, DC, T and B cell numbers

For G, 1 symbol represents 1 replicate, with the line showing median. For H-J, M, bars represent median, with each symbol representing 1 replicate. P values are for Mann-Whitney U test (*P<0,05, **P<0,01, ***P<0,001). Data are representative of 3 independent experiments. Related to figure 5.
